# Cancer modeling in colorectal organoids reveals intrinsic differences between oncogenic *RAS* and *BRAF* variants

**DOI:** 10.1101/860122

**Authors:** Jasmin B. Post, Nizar Hami, Jeroen Lohuis, Marieke van de Ven, Renske de Korte-Grimmerink, Christina Stangl, Ellen Stelloo, Ingrid Verlaan, Jacco van Rheenen, Hugo J.G. Snippert

## Abstract

Colorectal cancers (CRCs) with oncogenic mutations in *RAS* and *BRAF* are associated with anti-EGFR therapy resistance. Consequently, all RAS mutant CRC patients are being excluded from this therapy. However, heterogeneity in drug response has been reported between *RAS* mutant CRC patients. It is poorly understood to what extent such differences are derived from different genetic backgrounds or intrinsic differences between the various RAS pathway mutations. Therefore, using CRISPR technology we generated an isogenic panel of patient-derived CRC organoids with various RAS pathway mutations (i.e. *KRAS*^*G12D*^, *BRAF*^*V600E*^, *KRAS*^*G13D*^ and *NRAS*^*G12D*^). All RAS pathway mutants promote ERK activation and tumor growth. However, *KRAS*^*G12D*^ and *BRAF*^*V600E*^ mutations in particular conferred robust resistance to anti-EGFR therapy, both *in vitro* and *in vivo*. Moreover, untreated KRAS^G13D^ mutants showed fastest growth in mice but remained sensitive to anti-EGFR therapy. Together, introducing mutation-specific oncogene signaling in CRC organoids resembles clinical phenotypes and improves understanding of genotype-phenotype correlations.

## Introduction

The EGFR-RAS signaling pathway stimulates cellular proliferation during development, homeostasis and regeneration^1,2^. Consequently, aberrant activation of the signaling cascade by oncogenic mutations is frequently detected in cancers^3^. In metastatic colorectal cancer (mCRC), most abundant are mutations in RAS proteins or downstream kinase BRAF and to lesser extent at the receptor level^4,5^. KRAS, NRAS and HRAS are three RAS isoforms that propagate upstream EGFR signaling activity towards downstream RAF kinases^6^. Despite their ubiquitous expression pattern^7,8^ and high similarity in amino acid sequence (80%)^9–11^, the prevalence of oncogenic mutations across RAS isoforms are not equally distributed between different cancers^12^. Colorectal cancer is one of the most extreme cases, with mutations in *KRAS* being detected in 35-50% of the cases^13^, including even distribution across all tumorigenic stages^14^. In contrast, mutations in *NRAS* are only identified in 3-5% of mCRC patients and predominantly in malignant CRCs^13–15^, while HRAS mutations are virtual absent^12^. Moreover, the relative mutation frequency at hotspot locations differs between RAS isoforms, with G12 and G13 mutations most often detected in KRAS (combined 90%) and Q61 mutations frequently found in NRAS (57%)^16^.

These observations suggest that oncogenic variants of *RAS* isoforms are not equal during CRC development, including codon-specific influences. Indeed, mouse studies have demonstrated that the endogenous expression of *KRAS*^*G12D*^, but not *NRAS*^*G12D*^, promotes progression of intestinal neoplasia^17^. An increasing amount of observations seem at odds with the almost uniform classification of CRC tumors with a mutant EGFR signaling pathway. Accurate patient stratification based on RAS mutations is clinically relevant given that anti-EGFR targeted therapy has shown beneficial effects in mCRC patients without oncogenic RAS mutations^18,19^. However, just like oncogenic RAS variants being associated with different tumor phenotypes, various clinical studies demonstrated differences in therapeutic outcome for mCRC patients with different oncogenic RAS variants. For instance, they reported that mCRCs with *KRAS* codon 13 mutations are more often associated with a poorer prognosis compared to those with *KRAS* codon 12 mutations^20–22^. Paradoxically, patients with *KRAS* codon 13 mutations are reported to benefit from treatment with EGFR inhibitors^20–22^, in contrast to resistant *KRAS* codon 12 mutant tumors.

Unfortunately, the cause and consequences of potential differences between these oncogenic variants are poorly understood. In particular, intrinsic differences in tumorigenic potential between mutant variants is challenging to dissect, considering the immense different mutational backgrounds per human tumor^5^. Moreover, *NRAS* mutations are more often detected in tumors of the left colon, whereas *KRAS* mutant tumors are most frequently found in right colon^14,23^, creating the possibility that these are two different tumor subtypes of different epigenetic make-up and cellular composition. The situation is even more extreme for oncogenic mutations in *BRAF* (mostly V600E) that are detected in approximately 11% of CRC cases^4^. In contrast to RAS mutations, BRAF mutations are common in tumors with a hypermutation phenotype (microsatellite unstable)^24^, show co-occurrence with WNT pathway mutations in *RNF43* rather than in *APC*^4,5^ and show different metastatic behavior^16,23^.

To optimize personalized cancer treatment of CRC patients, we set out to improve our understanding of genotype-phenotype correlations and investigate the intrinsic similarities and differences between a series of common mutations in the EGFR signaling pathway. Patient-derived CRC organoids (CRC PDOs) are 3-dimensional ‘mini-organs’ that are established from primary tumor tissue, either obtained from biopsies or surgical resected material. CRC PDOs maintain the histopathological features of the native tumor, including high concordance between somatic mutations, transcriptome and drug response between matched primary tumors and derived organoid cultures^25–27^. Importantly, PDOs are compatible with CRISPR/Cas9-mediated introduction of cancer mutations^28,29^, and were used to generate a series of isogenic lines with various oncogenic RAS pathway mutations to reveal their similarities and differences.

## Results

### CRC PDOs with different MAPK pathway mutations display varying sensitivities to MAPK pathway inhibition

To document the intertumoral heterogeneity in drug response between CRCs with an oncogenic mutant MAPK signaling pathway, we performed a drug screen targeting various effectors in the MAPK pathway (**Fig. 1A and B**). Specifically, we applied this screen on a panel of CRC PDOs that are either wild-type or mutant for *KRAS, NRAS* or *BRAF* without displaying any other MAPK pathway mutations detected by Sanger sequencing (**Suppl. Fig. 1A**).

**Figure 1.**
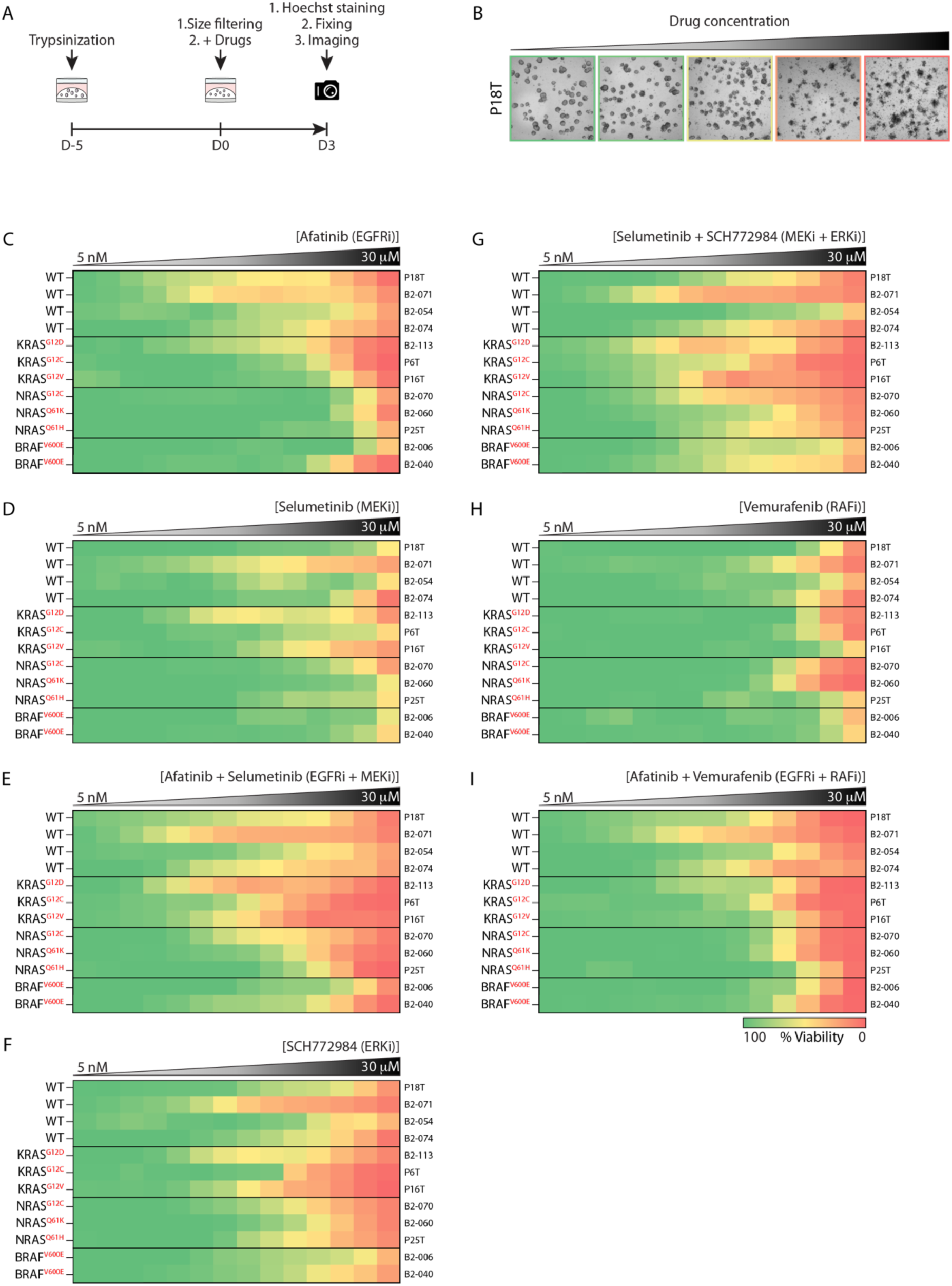
Differential drug sensitivities of *RAS* and *BRAF* mutant CRC PDOs to targeted MAPK pathway inhibition. (**A**) Schematic overview of the drug screening method. In short, a 3-day drug screen was initiated on 5 days old CRC PDOs expressing different *RAS* and *BRAF* mutations. (**B**) Representative bright field pictures of CRC PDOs (P18T) treated with increasing afatinib concentrations. (**C - I**) Heat maps of dose-response measurements (cell viability) in CRC PDO lines to (**C**) afatinib, (**D**) selumetinib, (**E**) afatinib plus selumetinib, (**F**) SCH772984, (**G**) SCH772984 plus selumetinib, (**H**) vemurafenib and (**I**) vemurafenib plus afatinib. Organoids were treated (72 hr) with vehicle (DMSO) or inhibitors targeting the EGFR-RAS-ERK pathway (5 nM – 30 μM range, in 14 logarithmic intervals). Red represents maximal cell death and green represents maximal viability. Drug names and their nominal targets are indicated above, the MAPK pathway mutant status per line at the left and the corresponding PDO line at the right. Average of 2 technical replicates.

In line with observations from the clinic, we observed that CRC PDOs with activating mutations in *RAS* and *BRAF* exhibited resistance to the pan-HER inhibitor afatinib^18,30,31^. The strongest sensitivity to pan-HER inhibition was observed in two out of the four *RAS*/*BRAF*^*WT*^ CRC PDOs (P18T and B2-071) (**Fig. 1C and Suppl. Fig. 1B**). We noticed limited response to pan-HER inhibition in the two remaining *RAS*/*BRAF*^*WT*^ CRC PDOs (B2-054 and B2-074). Potentially, these CRCs have MAPK pathway alterations that are either unknown, or were not identified by Sanger sequencing ^29,32–34^, but are in agreement with clinical observation of CRC patients enrolled in anti-EGFR therapy that show no response^35^.

In concordance with previous clinical and experimental observations, MAPK pathway inhibition downstream of RAS and BRAF with only the MEK inhibitor selumetinib was largely ineffective in the panel CRC PDOs^36–38^ (**Fig. 1D and Suppl. Fig. 1B**). However, upon combinatorial treatment with both pan-HER and MEK inhibitors, an improved drug response was predominantly detected in *K-* and *NRAS* mutant PDOs (**Fig. 1E and Suppl. Fig. 1B**).

Treatment with the ERK inhibitor SCH772984 showed the strongest effect on *KRAS* mutant and *RAS*/*BRAF*^*WT*^ CRC organoids, albeit at varying sensitivities (**Fig. 1F and Suppl. Fig. 1B**). Combinatorial treatment with ERK and MEK inhibitors induced an additive effect in most CRC organoids, irrespective of the mutational background, while variability in response between CRC PDOs remained largely unaffected (**Fig. 1G and Suppl. Fig. 1B**).

Response to BRAF inhibition with vemurafenib, largely ineffective in most CRC PDOs including organoids mutant for *BRAF*, corresponds with previous observations in CRC cell lines and clinical studies^39–42^ (**Fig. 1H and Suppl. Fig. 1B**). Strikingly, only a minor additive response was observed when pan-HER and BRAF inhibitors were co-administered in CRC PDOs (**Fig. 1I and Suppl. Fig. 1B**), recapitulating observations from the clinic about insufficient response rates in *BRAF* mutant patients^43^.

Remarkedly, differential drug sensitivities were observed between CRC PDOs with similarly affected oncogenes. In particular the *NRAS*^*G12C*^ mutant showed higher sensitivity to most drugs than the other *NRAS* mutants, either due to the type of point mutation or other tumor intrinsic properties. Similar type of deviation in drug response was observed between KRAS mutants, confirming previous reports on CRC organoid biobanks with various RAS mutants^26,44–46^. Overall, *BRAF* mutant CRC PDOs showed a very resistant phenotype to MAPK pathway inhibition. However, the nature of their primary tumors (e.g. sessile serrated adenomas) instead of their mutational status, could very well be responsible for the observed phenotype.

### Generation of endogenous oncogenic RAS and BRAF knock-in variants in CRC PDOs

To exclude the impact of genetic background, epigenetics and cellular composition on drug response when comparing various oncogenic mutations in the MAPK pathway, we set out to generate a panel of isogenic CRC PDOs harboring different <u>R</u>AS <u>p</u>athway <u>m</u>utations (RPMs). Apart from a non-functional APC and TP53 pathway, CRC PDO P18T has a genetically wild-type MAPK signaling pathway and shows sensitivity towards EGF-mediated tumor growth (**Fig. 1C and Suppl. Fig. 1B**)^25^. Therefore, we isolated CRC PDO P18T to introduce different oncogenic mutations in *KRAS, NRAS* and *BRAF* via homologous recombination (HR) using the CRISPR technology (**Fig. 2A**). Importantly, the donor template contained a puromycin selection cassette to enable selection independent of using the attributed mutant phenotype for selection like previously applied growth factor depletion regimens (e.g. EGF) (**Fig. 2A and 2B**)^47,48^. Next, we targeted CRISPR/Cas9 to the intronic region directly downstream of the exon of interest. Although this position is not optimal in relation to the distant upstream location of the point mutations, introducing a small intronic indel mutation in the non-recombined allele will have minimal to no impact on its integrity. Maintaining the expression of the wild-type allele is essential as it avoids allelic imbalance, a phenomenon that has been described during progression to malignancy where mutant KRAS gains the upper hand over wild-type^49–52^ (**Fig. 2A**).

**Figure 2.**
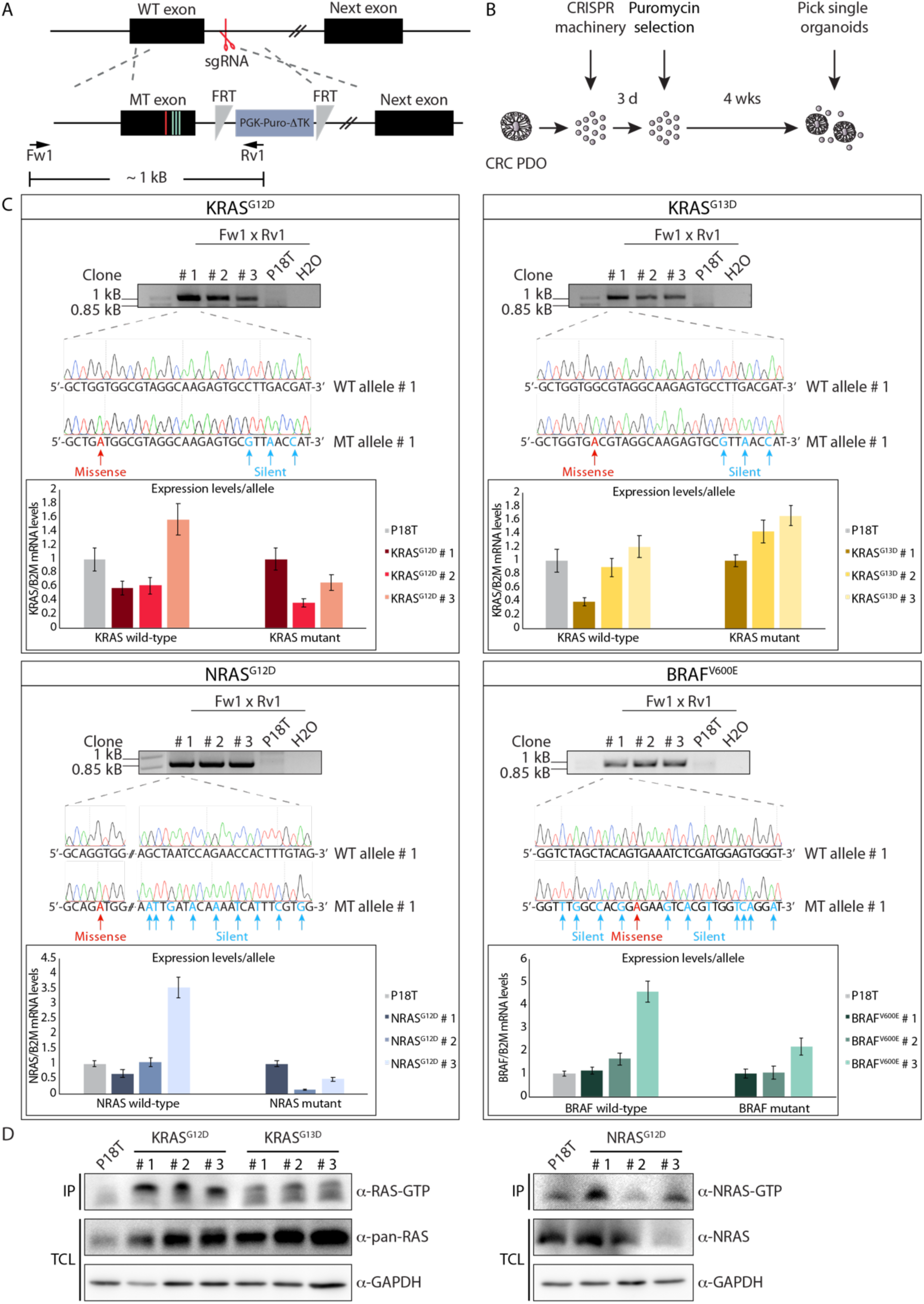
Generation of oncogenic *BRAF* and *RAS* knock-in variants in CRC PDOs. (**A**) Genetic strategy to target *KRAS, NRAS* and *BRAF* locus for homologous directed repair using CRISPR/Cas9 technology. Red and blue lines indicate oncogenic missense and silent mutations, respectively. Black boxes illustrate exons, separated by introns. Red scissor shows sgRNA-generated double stranded break. Black arrows illustrate PCR primer pairs that were used for the identification of knock-in clones. (**B**) Puromycin selection strategy to generate *RAS* and *BRAF* mutant CRC PDOs after CRISPR-mediated homologous recombination. (**C**) Per mutation the agarose electrophoresis gels showing the ∼1kb PCR product of the knock-in allele in 3 monoclonal lines per mutations. Sanger sequencing confirms presence of both knock-in and WT alleles. DNA sequences show introduction of missense (red) and silent (blue) mutations. The mRNA expression levels of wild-type and mutant alleles was analyzed using qPCR. The relative expression of each allele was normalized to the *B2M* housekeeping gene (representative from n = 3 independent experiments). (**D**) Western blot analysis shows enhanced RAS activity (GTP-loading) in *RAS* mutant CRC PDOs compared to P18T organoids. KRAS and NRAS immunoblots from RAS pull-down assays (RAS-GTP) and total lysates (loading control) are shown for *KRAS* and *NRAS* mutant organoids, respectively. Representative from n=2 independent experiments.

The generation of oncogenic mutations in *KRAS, NRAS* and *BRAF* were confirmed by DNA sequencing analysis of three independent monoclonal lines (**Fig. 2C**), confirming the presence of both mutant and wild-type alleles. Additional DNA sequence analyses of the intronic regions targeted by Cas9 confirmed the generation of indels in the allele that was not subjected to HR (**Suppl. Fig 2A and 2B**). In agreement, both wild-type and mutant alleles were shown to be expressed at near equal ratios in the isogenic RPM lines as demonstrated by qPCR analysis using allele-specific primers (**Fig. 2C**). Subsequent RAS-GTP pull down assays with the RAS binding domain (RBD) of RAF confirmed presence of oncogenic RAS proteins, including overall correlation between increased active RAS-GTP loading levels and mRNA expression of the mutant allele (**Fig. 2D**). Moreover, we observed increased protein levels of total RAS in *RAS* mutant organoids, indicative of positive feedback regulation induced by oncogenic *RAS* proteins as previously reported^53–55^.

### BRAF^V600E^ and KRAS^G12D^ knock-in CRC PDOs show resistance to pan-HER inhibition

To maintain clonal diversity, we continued with three independent clones per oncogenic mutation and investigated their effect on EGF-dependent tumor growth.

Therefore, we monitored organoid number, size, and viability at different time points after organoid plating and drug administration (**Suppl. Fig 3A**). Under normal growth conditions, we observed no significant differences between parental P18T and oncogenic RPM lines (**Fig. 3A**). However, significant differences were observed when we challenged the RPM organoids with the pan-HER inhibitor afatinib to block all EGF-stimulated receptor signaling. Whereas all oncogenic RPM organoids had a survival benefit compared to parental P18T organoids upon short-term (3 days) afatinib treatment, organoids expressing *KRAS*^*G12D*^ and *BRAF*^*V600E*^ oncogenes showed most resistance to afatinib that became most prevalent during long-term (7 days) treatment (**Fig. 3B and Suppl Fig. 3B**). In particular, relapse after drug treatment showed a striking growth of both *KRAS*^*G12D*^ and *BRAF*^*V600E*^ mutant organoids, while the others were significantly depleted in number or showed growth arrest (**Fig. 3C-D and Suppl. Fig. 3C**). While overall phenotypes per mutation showed resemblance, we did observe interclonal heterogeneity, in particular for NRAS. Presumably epigenetic and/or transcriptional differences between the cells-of-origin of the mutant clones can still affect phenotype^26,45^. For example, the most resistant NRAS^G12D^ clone (#1) correlates with highest expression level and GTP-loading of the mutant protein (**Fig. 2C and 2D**).

**Figure 3.**
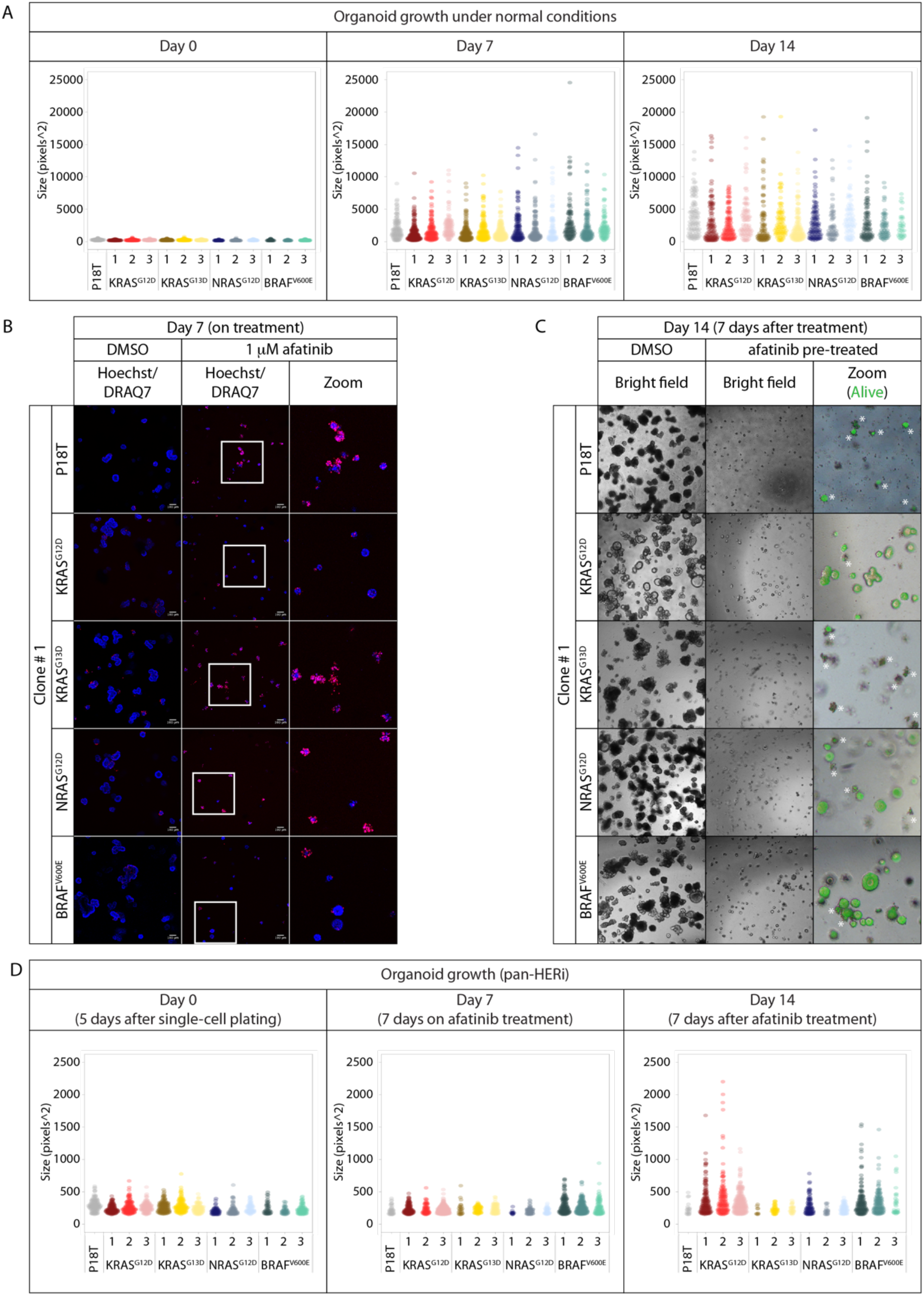
*KRAS*^*G12D*^ and *BRAF*^*V600E*^ knock-in mutations promote organoid survival and growth. (**A**) Quantitative analysis of organoid size and number in isogenic RPM lines during normal culture conditions (DMSO) at different time points after organoid plating. Size and number of viable organoids were measured by uptake of fluorescent calcein green (see methods). Each dot represents one organoid^76^. Data from 1 8-well Lab-Tek chambered coverglass is shown. (**B**) Representative fluorescent pictures of parental P18T, or *KRAS*^*G12D*^, *KRAS*^*G13D*^, *NRAS*^*G12D*^ and *BRAF*^*V600E*^ knock-in organoids (clone # 1) after 7 days of DMSO or afatinib treatment (1 μM). Scale bars, 100 μM. Hoechst (blue) and DRAQ7 (red) was used to visualize nuclei and dead cells, respectively. (**C**) Representative bright field pictures of parental P18T, or *KRAS*^*G12D*^, *KRAS*^*G13D*^, *NRAS*^*G12D*^ and *BRAF*^*V600E*^ knock-in organoids (clone # 1) 7 days after release of afatinib treatment (day 14). Representative zoom-in panels show fluorescent calcein green signal in living cells. Asterisks indicate autofluorescence of dead material (**Suppl. Fig 3C**). (**D**) Quantitative analysis of organoid growth and viability in isogenic RPM lines prior (day 0), during (day 7) and after (day 14) treatment with afatinib (1 μM). Size and number of viable organoids were measured by uptake of fluorescent calcein green (see methods). Each dot represents one organoid^76^. Data from 1 8-well Lab-Tek chambered coverglass is shown.

Remarkedly, regardless of oncogenic RAS pathway mutations, we observed that growth remains dependent on additional HER-mediated signaling input (**Fig. 3B and D**). Although the HER receptor family is upregulated in the RPM organoids (**Suppl. Fig. 3D**), we noticed that EGFR signaling was mostly responsible for this (**Suppl. Fig. 3E**).

### BRAF^V600E^ and KRAS^G12D^ knock-in CRC PDOs show residual MAPK pathway activation upon pan-HER inhibition

Previous studies have indicated that the lack of sensitivity towards MAPK pathway inhibition in *RAS* and *BRAF* mutant CRCs is caused by residual levels of ERK activity^41,56,57^. To investigate the molecular mechanism underlying resistance to pan-HER inhibition as observed in *KRAS*^*G12D*^ and *BRAF*^*V600E*^ RPM organoids, we analyzed MAPK signaling activity using biochemistry. As expected, introduction of the oncogenic mutations in RPM organoids resulted in increased ERK activation under normal culture conditions (**Fig. 4A, Suppl. Fig. 4A**). In line with the sensitive phenotypes during pan-HER inhibition, we noticed complete loss of MEK and ERK activity in P18T organoids as well as in *KRAS*^*G13D*^ and *NRAS*^*G12D*^ RPM organoids. In contrast, residual MEK and ERK activity was observed in RPM organoids expressing *KRAS*^*G12D*^ and *BRAF*^*V600E*^, albeit with different kinetics (**Fig. 4A, Suppl. Fig. 4A**). Initially, all organoids with oncogenic *RAS* variants displayed a fast and full inhibition of ERK phosphorylation upon pan-HER inhibition. Subsequently, over the course of 24 hours most reactivation of ERK was observed in the presence of mutant KRAS^G12D^. Unlike oncogenic *RAS* variants, the drug response of *BRAF*^*V600E*^ mutant organoids towards pan-HER inhibition displayed slower kinetics of downstream pathway inhibition (**Fig. 4A, Suppl. Fig. 4A**). Moreover, even after 72 hours we noticed no strong reactivation of ERK as consistently observed in *KRAS*^*G12D*^ organoids, but minimal levels that remained stable over time (**Suppl. Fig. 4B**). Notably, the minimal levels of ERK phosphorylation observed after 24 hours of pan-HER inhibition in *BRAF*^*V600E*^ mutants seem comparable to *KRAS*^*G13D*^ organoids (**Fig. 4A, Suppl. Fig. 4A**), but their sensitivity towards pan-HER inhibition deviates substantially (**Fig. 3B-D**). This suggests that minimal differences in ERK activity at the lower range can induce different cellular effects.

**Figure 4.**
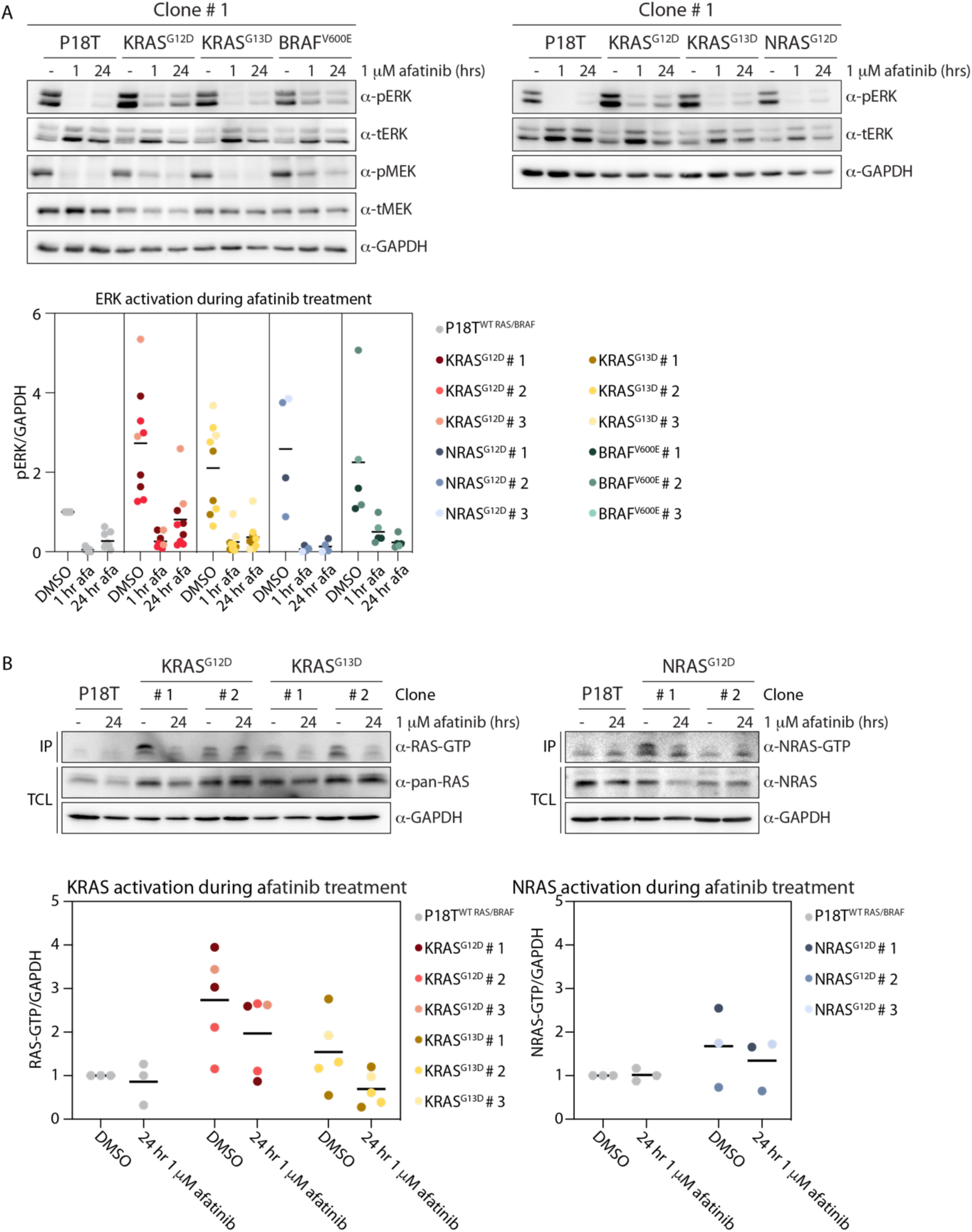
*KRAS*^*G12D*^ and *BRAF*^*V600E*^ RPM lines show residual MAPK pathway activity in the presence of pan-HER inhibition. (**A**) Organoids expressing oncogenic *KRAS* (G12D and G13D), *NRAS* (G12D) and *BRAF* (V600E) variants show enhanced basal ERK phosphorylation levels compared to P18T organoids. Pan-HER inhibition (1 µM afatinib) shows sustained ERK and MEK phosphorylation in *KRAS*^*G12D*^ and *BRAF*^*V600E*^ organoids compared to P18T, *KRAS*^*G13D*^ and *NRAS*^*G12D*^ organoids. Top panels are representative biochemistry experiments on clone # 1 from n=3. Bottom scatter plot depicts ERK phosphorylation levels normalized to GAPDH for all clones (n≥4). Baseline of P18T (DMSO) is set at 1. (**B**) Top panels depict biochemistry on RAS activity (GTP-loading) in unperturbed culture conditions and pan-HER inhibition (1 µM afatinib) for *KRAS* (G12D and G13D) and *NRAS* (G12D) mutant clones # 1 and 2 compared to P18T CRC organoids. RAS immunoblots from RAS pull-down assay are shown (RAS-GTP), together with a RAS immunoblot from total cell lysates as loading control. HRAS, KRAS, and NRAS isoforms are detected in mutant KRAS pull-down assays. NRAS isoforms are detected in mutant NRAS pull-down assays. Representative from n = 3 independent experiments. Scatter plots below depict RAS-GTP levels normalized to GAPDH for all clones (n≥3). Baseline of P18T (DMSO) is set at 1.

The level of active RAS-GTP was elevated in almost all *RAS* mutant RPM organoids in unperturbed culture conditions (**Fig. 4B and Suppl. Fig. 4C-D**) with the exception of *NRAS*^*G12D*^ # 2, which is consistent with the limited expression levels of the mutant allele. Subsequently, an overall decrease in RAS-GTP loading was observed for all mutant lines in the absence of upstream EGFR signaling input, although the degree of reduction varied per mutation (least in *KRAS*^*G12D*^, most in *KRAS*^*G13D*^) and per clone.

### Interclonal variability on karyotype and transcriptome

All RPM clones were created from the same polyclonal P18T organoid culture. As a result, clonal RPM lines can originate from different subclones present in the parental culture and might underly interclonal variability. To interrogate the clonal origin of the RPM lines, we determined DNA copy number alterations (CNAs) of the RPM organoids to determine clonal-specific karyotypes. Overall, a high degree of similarity in the pattern of CNAs was observed across all isogenic RPM organoid lines and the parental P18T organoids (**Fig. 5A**). Moreover, amplification of the *BRAF* allele as indicated in the CNA plots, corresponds with the detection of two wild-type alleles with independent Cas9-created indels in BRAF^V600E^ # 1 (**Suppl. Fig. 2B**). Notably, RPM clones with the highest passage numbers (clone #1, passage number data not shown) showed most extreme deviations from the average karyotype, indicative of ongoing clonal evolution during culture as previously reported^26^. In addition to ongoing chromosomal instability, certain clones displayed a higher degree of CNA resemblance than to others. In particular, a gain of chromosome 2 in half of the clones and (sub)chromosomal loss of chromosome 4 in most but not all. The observed gross similarities in karyotypes is likely the result of monoclonal lines originating from different clonal populations that were present in the polyclonal bulk P18T culture. In particular, NRAS^G12D^ # 1 that is the most phenotypic outlier of the NRAS^G12D^ mutants also displays most genomic alterations, potentially supporting the concept that clonal origin may underly interclonal variability. Nevertheless, overall phenotypic response towards EGF-independent growth conditions seems primarily dictated by the specific RAS pathway mutation and independent of clonal deviations.

**Figure 5.**
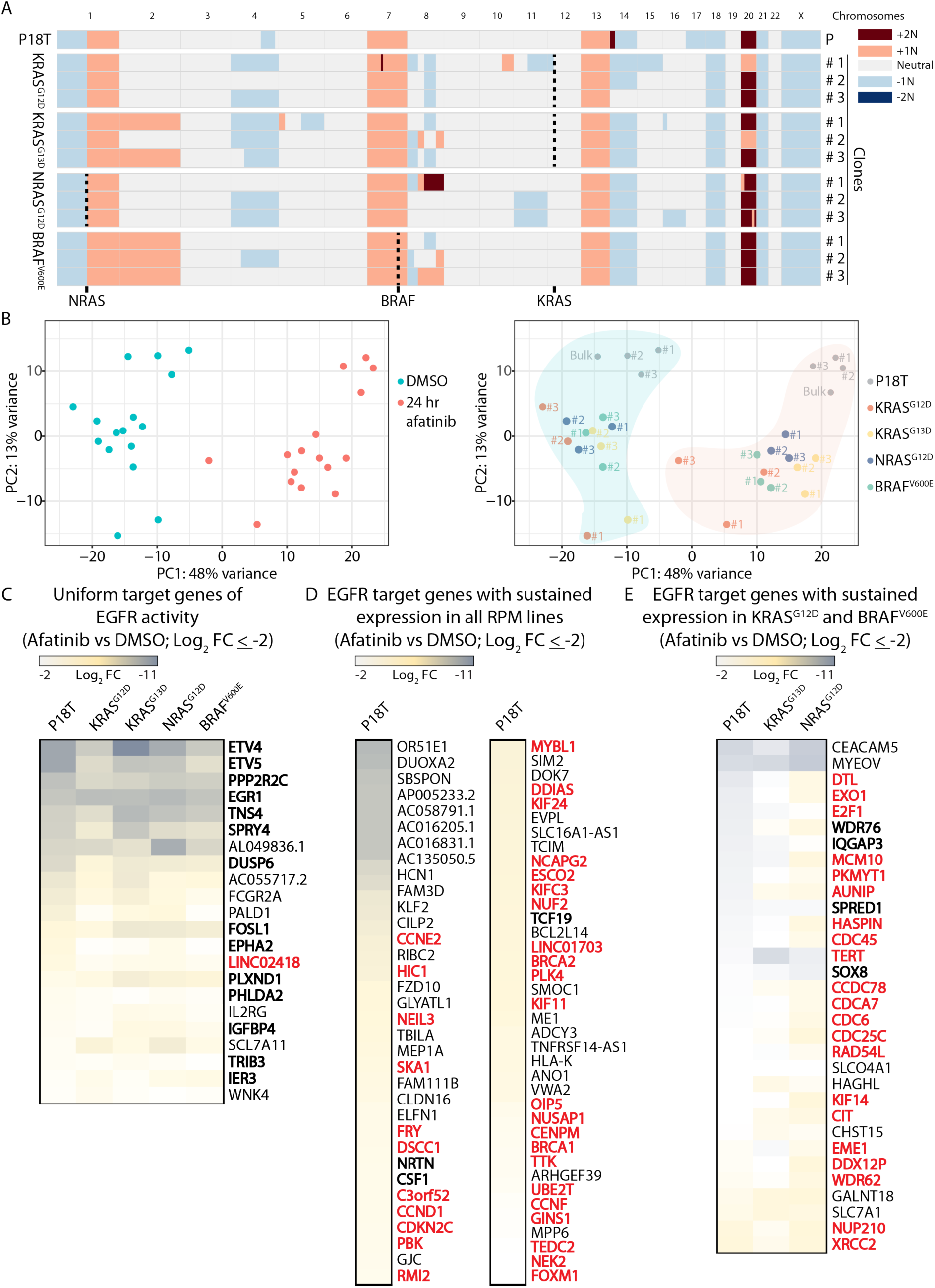
Karyotype and gene expression profiles of RPM lines. (**A**) DNA copy number alterations of monoclonal RPM lines (clones # 1-3) and parental (P) P18T organoids. Centromeric regions (thin black line in chromosomes) are excluded due to repetitive nature of DNA sequence. Dashed lines indicate the genetic location of *BRAF, KRAS* and *NRAS loci*. Colors per (sub)chromosome indicate CNA according to legend. (**B**) Principle Component Analysis of gene expression levels for monoclonal RPM lines and parental P18T organoids (bulk and monoclonal lines # 1-3) during unperturbed growth conditions (DMSO) and pan-HER inhibition (1 µM afatinib for 24 hr). Left panel indicates treatment. Right panel indicates clone identities. Hue at background corresponds to treatment. (**C**) Heatmap of significantly downregulated genes (p_-adjusted_ <0.05; log_2_ fold change (FC) ≤-2) after afatinib treatment in all RPM lines and parental P18T organoids gives uniform EGFR target genes (**D**) As in (C), but EGFR signature genes as determined in P18T organoids that sustain expression levels in all RPM lines. (**E**) As in (D), but EGFR signature genes as determined in P18T organoids with only sustained expression levels in resistant *KRAS*^*G12D*^ and *BRAF*^*V600E*^ clones. RAS-ERK pathway related genes are depicted in bold. Cell cycle-related genes are marked in bold, red font.

Next, we performed RNA sequencing analysis to investigate the degree of clonal variability on the transcriptional level, both during normal growth conditions as well as during pan-HER targeted therapy. To measure the variance in global gene expression between all RPM lines, we performed Principle Component Analysis (PCA, Methods). Based on the first two principle components, the largest differences in gene expression were observed between DMSO and afatinib treated conditions and between wild-type and mutant RAS pathway organoids (**Fig. 5B**). Moreover, during normal growth conditions we noticed somewhat random clustering between all RPM clones, but a tendency to cluster per mutation/phenotype during EGF-deprived signaling with KRAS^G12D^ and BRAF^V600E^ clustering towards the center of the PC1 axis. Subsequently, we determined the transcriptional signature mediated by EGFR signaling in CRC organoids in relation to RAS/RAF oncogene-mediated transcriptional activity. Using the parental P18T organoids with a wild-type MAPK pathway, we identified the EGFR gene signature by perturbing receptor activity using pan-HER inhibition (266 differentially expressed genes) (**Suppl. Fig. 5A and Suppl. Table 1**). Intriguingly, this EGFR gene signature with many known targets of the MAPK pathway remains to a large extent dependent on upstream EGFR signaling activity in the RPM mutants (**Fig. 5C, Suppl. Fig. 5A and Suppl. Table 1**), in agreement with the significant reduction in ERK activity upon afatinib treatment. Target genes of the EGFR signature that are least affected in the RPM mutants relate to cell cycle pathways, which is most dramatic for KRAS^G12D^ and BRAF^V600E^ RPM lines and in line with the observed phenotypes (**Fig. 5D-E, Suppl. Fig. 5B and Suppl. Tables 1-3**). Moreover, when directly comparing differential gene expression between afatinib-resistant (i.e. KRAS^G12D^ and BRAF^V600E^) and -sensitive clones (i.e. KRAS^G13D^ and NRAS^G12D^) independently analyzed from the EGFR gene signature, we again observed lower expression of cell cycle-related genes in sensitive organoids during treatment (**Suppl. Fig. 5B-C**).

### Intertumoral heterogeneity between MAPK oncogenic mutations during drug responses

To improve the characterization of drug response phenotypes in relation to KRAS, NRAS and BRAF oncogenes, we performed a drug screen on the RPM organoids with various targeted inhibitors against the MAPK pathway (**Fig. 6A**).

**Figure 6.**
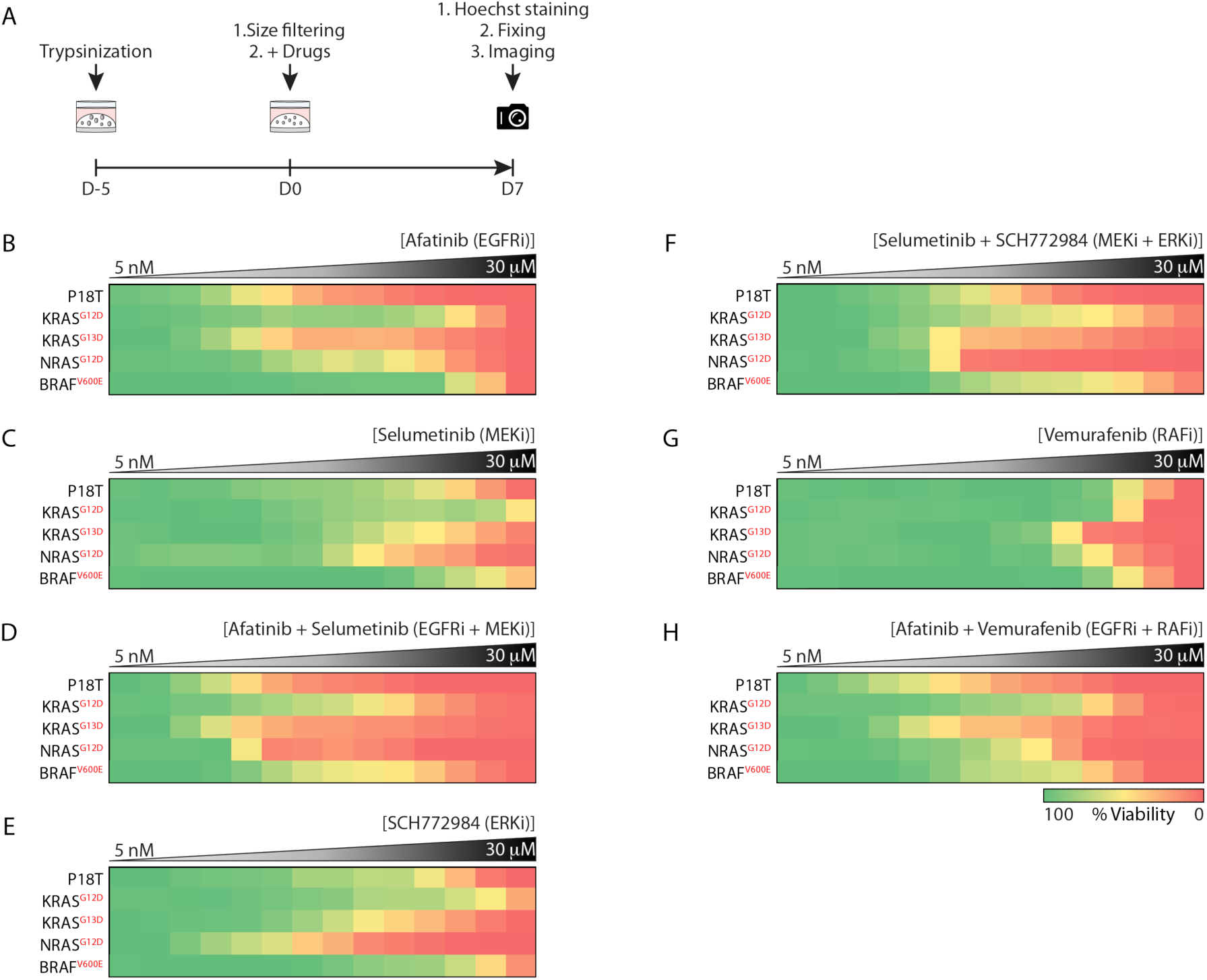
Differential drug sensitivities of oncogenic *RAS* and *BRAF* knock-in CRC PDOs to targeted MAPK pathway inhibition. (**A**) Schematic overview of the drug screening method. In short, a 7-day drug screen was initiated on 5 days old P18T and RPM organoids expressing different *RAS* and *BRAF* mutations. (**B - H**) Heat maps of dose-response measurements (cell viability) in CRC PDO lines to (**B**) afatinib, (**C**) selumetinib, (**D**) afatinib plus selumetinib, (**E**) SCH772984, (**F**) SCH772984 plus selumetinib, (**G**) vemurafenib and (**H**) vemurafenib plus afatinib. Organoids were treated (7 days) with vehicle (DMSO) or inhibitors targeting the EGFR-RAS-ERK pathway (5 nM – 30 μM range, in 14 logarithmic intervals). Red represents maximal cell death and green represents maximal viability. Drug names and their nominal targets are indicated above and the MAPK pathway mutant status per line at the left. Average of 2 technical replicates.

Confirming previous observations, *KRAS*^*G12D*^ and *BRAF*^*V600E*^ RPM organoids showed resistance to pan-HER inhibition with afatinib, with NRAS^G12D^ mutants being an intermediate and KRAS^G13D^ phenocopying the parental P18T line (**Fig. 6B and Suppl. Fig. 6A**).

While MEK inhibition alone showed limited effect, intertumoral variability was as expected (**Fig. 6C and Suppl. Fig. 6A**). Moreover, combined inhibition of MEK and pan-HER showed high sensitivity for P18T, *KRAS*^*G13D*^ and *NRAS*^*G12D*^ mutant organoids, with a pronounced additive effect on *NRAS*^*G12D*^ organoids (**Fig. 6D and Suppl. Fig. 6A**). Monotherapy against ERK with SCH772984 showed a similar pattern as MEK inhibition, albeit with overall lower IC50s (**Fig. 6E and Suppl. Fig. 6A**), while intertumoral variability in drug response upon combining MEK and ERK inhibitors resembled pathway inhibition via pan-HER and MEK inhibition (**Fig. 6F and Suppl. Fig. 6A**). General insensitivity towards BRAF inhibition with vemurafenib was observed in all RPM organoids, including BRAF^V600E^ (**Fig. 6G and Suppl. Fig. 6A**), with minimal additive effects upon its combination with dominant pan-HER inhibition (**Fig. 6H and Suppl. Fig. 6A**). Together, the data confirm earlier observations that KRAS^G12D^ and BRAF^V600E^ impose a more resistant phenotype towards EGF-independent growth conditions than KRAS^G13D^ and NRAS^G12D^. In addition, unique targetable vulnerabilities per mutation type were not identified within the MAPK pathway.

Next, we decided to focus on multiple patient-derived BRAF^V600E^ mutants (CRC PDOs and BRAF^V600E^ RPMs) in order to keep the oncogenic mutation a constant, while varying between separate tumors. The 3-day drug response phenotypes of *BRAF*^*V600E*^ RPM lines showed overall high resemblance to the other *BRAF* mutant CRC PDOs. Yet, intertumoral heterogeneity was observed between the lines where B2-040 showed most sensitive phenotypes. This confirms the widely supported notion that differences beyond the oncogenic mutation, among others differences in (epi)genetic landscape, cellular composition and transcriptional levels, will additionally impact tumor cell intrinsic drug sensitivities (**Suppl. Fig. 6B-I**).

### In vivo tumor growth and response to anti-EGFR therapy

To characterize the similarities and differences of the RPM lines in vivo, we xenografted the RPM organoids in mice to analyze growth dynamics (**Fig. 7A**). As expected, tumors from isogenic RPM organoids displayed faster growth rates as compared to the parental P18T organoids (**Fig. 7B**). Moreover, we observed that *KRAS*^*G13D*^ mutant tumors showed the fastest growth rates compared to other RPM tumors, resembling elevated tumor growth dynamics of mCRC patients with *KRAS*^*G13D*^ mutations^20–22^. More surprising to us was that KRAS^G12D^ showed slowest growth kinetics of the RPM lines, since endogenous expression of KRAS^G12D^, but not NRAS^G12D^, promotes cancer progression in murine colonic epithelium after loss of APC expression^17^.

**Figure 7.**
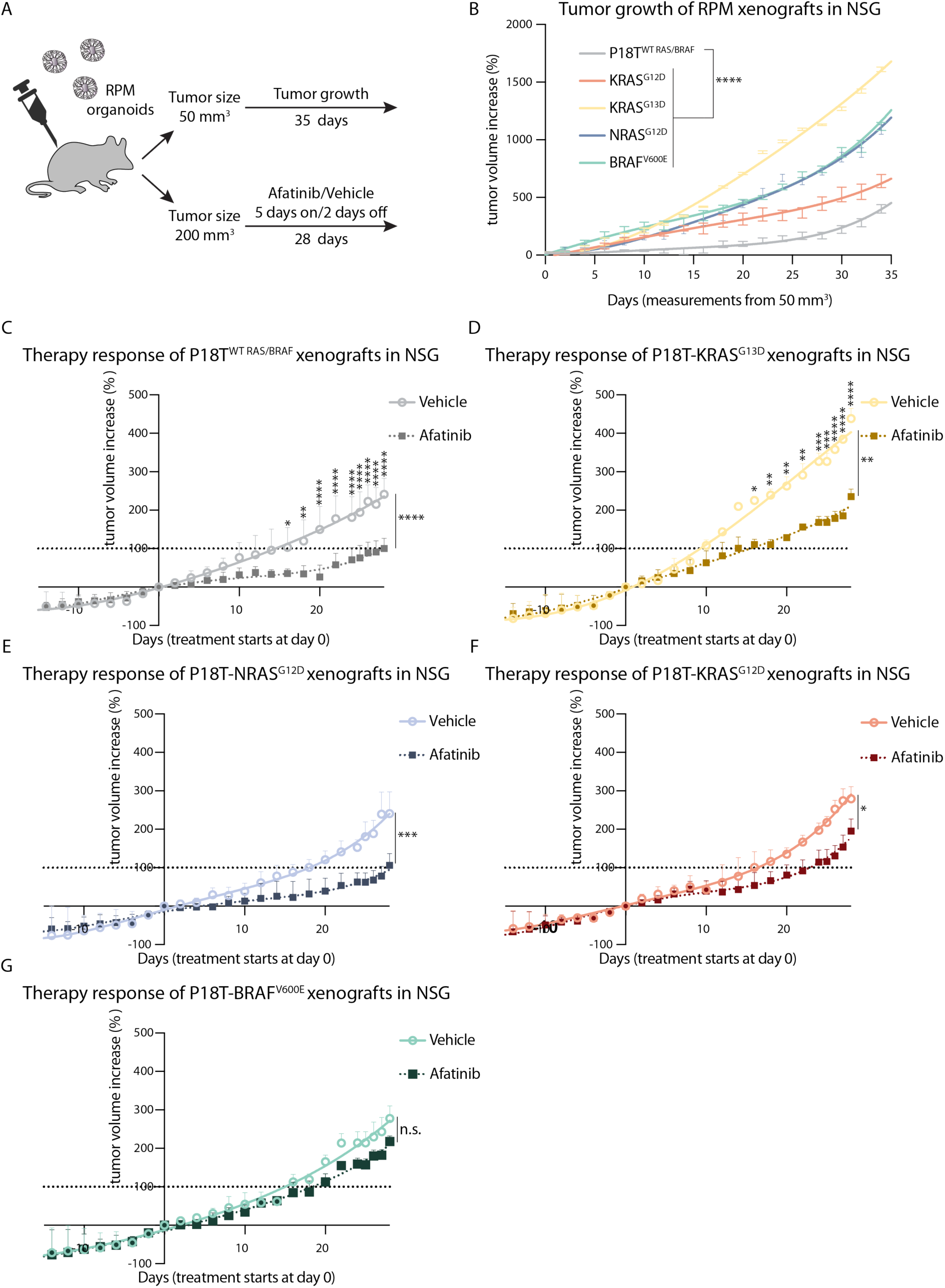
In vivo growth dynamics of RPM organoids. (**A**) A schematic overview illustrating in vivo examination of xenografted RAS pathway mutant (RPM) CRC PDOs in mice. In short, CRC PDOs were subcutaneously injected in mice. Above 50 mm^3^, growth of tumors was analyzed for 35 days. Above 200 mm^3^, tumors were treated with either vehicle or afatinib (20 mg/kg via oral gavage, 5 days on/2 days off) for 28 days. (**B**) Graph displays the mean percentage change in tumor volume relative to initial tumor volume (50 mm^3^) of xenografts from indicated RPM organoids monitored over the experimental period. Error bars represent standard deviation. n = 5 per group. **** p < 0.0001, 2-way ANOVA Dunnett’s multiple comparisons test. (**C**) Graphs display the mean percentage change in tumor volume relative to initial tumor volume (200 mm^3^) of xenografts from P18T treated with vehicle or afatinib. (**D**) like (C) but for KRAS^G13D^ (**E**) NRAS^G12D^, (**F**) KRAS^G12D^ and (**G**) BRAF^V600E^. Error bars represent standard deviation. n = 5 per group. p > 0.05; * p < 0.05; ** p < 0.01; **** p < 0.0001; n.s., not significant. For multiple comparison per time point 2-way ANOVA (Bonferroni’s multiple comparisons test) was used. For overall *in vivo* response Welch’s *t*-test was used.

Multiple studies have been published reporting the accuracy of tumor-derived organoids in predicting patient responses to therapy^27,58,59^. To verify whether our observed intertumoral differences in drug responses between the RPM lines are also observed *in vivo*, we applied afatinib treatment for 28 days. Consistent with results from *in vitro* experiments, a significant decrease in growth rate was observed in P18T, *KRAS*^*G13D*^ and *NRAS*^*G12D*^ mutant tumors upon afatinib treatment (**Fig. 7C-E**). In comparison, *BRAF*^*V600E*^ and *KRAS*^*G12D*^ mutant tumors were most resistant, showing only a marginal reduction in tumor growth compared to vehicle (**Fig. 7 F-G**). Although overall *in vitro* responses were more dramatic, presumably due to a mismatch between achieved drug concentrations *in vivo*, the overall pattern of intertumoral differences between the RPM clones remained present.

*KRAS*^*G13D*^ mutant tumors are an interesting outlier, considering its malignant nature during unperturbed growth conditions in mice. Yet, sensitivity to anti-EGFR therapy remains, so suppressed growth rates could be established towards the range of vehicle-treated parental P18T tumors (**Fig. 7D**). This corresponds to observations from the clinic, showing that patients with *KRAS*^*G13D*^ mutant CRCs benefit from cetuximab treatment, but are inferior to the response of CRC patients with *RAS*^*WT*^ tumors^20,22,60^.

## Discussion

Despite the overlapping functions of RAS and BRAF oncogenes, data analyses of CRC patients and cell lines suggest that mutation variants of oncogenic RAS and BRAF proteins are associated with different tumor types and involved in distinct tumorigenic pathways^17,23,61,62^. Moreover, CRCs with oncogenic *RAS* variants can react differently to pan-HER inhibition^20–22,63,64^. Nevertheless, stratification of patient cohorts per mutation is significantly hampered by limited statistical power due to small sample sizes. Therefore, mCRC patients with any activating mutation in *RAS* are currently being excluded from anti-EGFR targeted therapy^65^. Moreover, from a practical point of view it is very challenging to dissect similarities and differences between various oncogenic RAS and BRAF variants using patient-derived material. Foremost, because intertumoral heterogeneity is to a large extent influenced by differences in genetic landscape of the tumor, epigenetics, cellular composition, tumor location, stroma infiltration, and more^4,23,24,62,66^.

Therefore, to address the influence of a single RAS pathway mutation on tumor progression and anti-EGFR therapy resistance, we generated a panel of isogenic CRC PDO lines with endogenous expression of a variety of oncogenic *BRAF* and *RAS* mutations. Confirming the oncogenic role of MAPK pathway mutations, we observed that activating mutations in *BRAF* (i.e. V600E), *KRAS* (i.e. G12D and G13D) and *NRAS* (i.e. G12D) enhanced downstream MAPK pathway activity. In the presence of exogenous EGF levels during normal culture conditions, growth analysis and RNA sequencing did not reveal significant differences between the RPM lines. In contrast, under more challenging conditions as EGF-independent growth due to pan-HER inhibition, we noticed apparent differences between the RPM mutants where KRAS^G12D^ and BRAF^V600E^ showed most resistant phenotypes. Same differential sensitivity towards pan-HER inhibition was also observed *in vivo*.

During normal growth conditions *in vivo* where EGF supply is limited but not absent, *KRAS*^*G13D*^ RPM lines displayed enhanced tumor growth over the other RPM tumors. This corresponds to the malignant nature of *KRAS*^*G13D*^ mutant CRCs in patients that is likely caused by the high level of intrinsic GDP-to-GTP exchange rates of KRAS^G13D^ mutant proteins, resulting in guanine nucleotide exchange factor (GEF) independent RAS auto-activation^20,22,60,67,68^. The codon-specific effects of KRAS^G13D^ mutants are likely masked in *ex vivo* culture conditions with EGF concentrations that saturate EGFR phosphorylation and downstream pathway activation. The structural differences between KRAS^G12D^ and KRAS^G13D^ proteins might explain their distinct response to pan-HER inhibition^68,69^, as KRAS^G13D^, but not KRAS^G12D^, retain sensitivity to GAP-mediated GTP hydrolysis as demonstrated by *in vitro* GTP hydrolysis experiments^67,68^.

The high prevalence of codon 12 and 61 mutations in *KRAS* and *NRAS*, respectively, suggests that the transforming capacity differs per oncogenic mutation and RAS isoform^6^ (COSMIC database). We observed that *NRAS*^*G12D*^ were ‘weaker’ mutants than *KRAS*^*G12D*^ in terms of EGF-independent growth phenotypes of our RPM lines. In contrast, a NRAS^Q61^ mutant CRC PDO showed most resemblance to KRAS^G12D^. It has been reported that *NRAS* codon 61 mutants are more efficient inducers of downstream MAPK signaling than *NRAS* codon 12^70^. We observed fewer active RAS proteins in NRAS^G12D^ RPM organoids compared to KRAS^G12D^ RPM organoids. A possible explanation for the preference of codon 61 mutations over codon 12 in *NRAS* is that lower expression levels require compensation by a potent mutation in order to achieve sufficient MAPK activation and transformative capacity. For *KRAS*, codon 61 mutations might create too much oncogenic potency, as KRAS^Q61^ mutations appear more active than KRAS^G12D^ mutations *in vitro*, and result in growth arrest of primary lung fibroblast^50,71^.

Although *KRAS*^*G12D*^ and *BRAF*^*V600E*^ RPM organoids showed resistance to pan-HER inhibition, a strong reduction in ERK activation and cell proliferation was observed. This indicates that upstream receptor activation is important to elevate levels of ERK activity, despite presence of MAPK pathway activating mutations. Conceptually, this parallels other observations in *RAS* mutant cancer cells, showing that the capacity of RAS and RAF oncogenes to activate downstream MAPK signaling is not maximal^72,73^. Moreover, the residual levels of RAS-ERK activity in *RAS* mutant organoids during drug screens did correlate with observed phenotypes, where ERK phosphorylation levels were significantly higher in afatinib-resistant (KRAS^G12D^) clones compared to afatinib-sensitive (KRAS^G13D^ and NRAS^G12D^) clones. In contrast, *BRAF*^*V600E*^ mutant clones showed a different drug response with respect to ERK phosphorylation than the oncogenic RAS variants, with slower kinetics and an incomplete inhibition but stable suppression over time. Considering that both the *KRAS*^*G12D*^ and *BRAF*^*V600E*^ mutant organoids manifest similar resistance phenotypes, we speculate that both drug response kinetics, i.e. either suppressed and sustained over time or initially eliminated with subsequent minimal reactivation, effectuate low ERK activity levels that are sufficient to support organoid survival, but not growth. Intriguing, while the minimal levels of ERK activity in *BRAF*^*V600E*^ and *KRAS*^*G13D*^ mutant organoids seem comparable, their phenotypes deviate significantly. Most likely, minimal differences in ERK activity levels at the lower range might have different phenotypic outcomes. Future methodologies for sensitive quantifications of signaling activities at the single cell level and in real-time will prove instrumental for full understanding how pathway activity levels in tumors correlate with drug response phenotypes (manuscript in preparation), among others to discriminate between heterogeneities of signaling activities between cells and fluctuating activity levels over time.

Together, we show that oncogenic knock-in mutations in the MAPK pathway in CRC PDOs resemble phenotypes observed in the clinic and can be used to understand mutation-specific oncogene signaling in CRC. Furthermore, we observed differential sensitivities to MAPK pathway inhibition between RAS pathway mutations, supportive of the notion that mCRC patients with *NRAS*^*G12D*^ and *KRAS*^*G13D*^ mutations, which are currently being excluded from anti-EGFR targeted therapy, might actually benefit from anti-EGFR therapy. Further investigation will be required to determine whether the current stratification of mCRC patients based on overall *RAS* mutant status should be reconsidered and include codon- and isoform-specific variations. Indeed, in addition to intrinsic differences between various mutations as revealed with the RPM organoids, cross-comparison of different CRC PDOs with identical BRAF^V600E^ mutation displayed intertumoral heterogeneity beyond the exact mutation type, like genetic landscape, epigenetics, cellular composition, tumor location and stroma infiltration. How all these multiple variables, including mutation type, act in concert in establishing a (drug response) phenotype is challenging to dissect as tumors have often adapted in multiple possible scenarios to accomplice sufficient MAPK activity. Future decision-making for therapeutic strategy should use the most optimal genotype-phenotype correlations to assist in overall patient stratification to narrow down on potential therapeutic options, after which tailored-made adjustments can be made based on personalized drug response data.

## Materials and methods

### Patient-derived organoid culture and maintenance

The patient-derived P6T, P16T, P18T and P25T organoids used in this study were previously established and characterized^25^. Other patient-derived organoids described in this study were established and characterized by the Hubrecht Organoid Technology (hub4organoids.eu). Human CRC organoids were cultured as described previously^25,44^. Culture medium contained advanced DMEM/F12 medium (Invitrogen) with 1% Penicillin/Streptomycin (P/S, Lonza), 1% Hepes buffer (Invitrogen) and 1% Glutamax (Invitrogen), 10% R-spondin conditioned medium, 10% Noggin conditioned medium, 1x B27 (Invitrogen), 1.25 mM n-Acetyl Cysteine (Sigma-Aldrich), 10 mM Nicotinamide (Sigma-Aldrich), 50 ng/ml EGF (Invitrogen), 500 nM A83-01 (Tocris), 10 μM SB202190 (ApexBio) and 100 μg/ml Primorcin (Invitrogen). Organoids were splitted through Trypsin-EDTA (Sigma-Aldrich) treatment. Culture medium after splitting was supplemented with 10 μM Y-27632 dihydrochloride. For selection of RAS pathway mutants, organoids were grown in culture medium containing 1-2 μM puromycin.

### Organoid transfection and genotyping

The transfection protocol of P18T organoids was previously described in detail by Fujii *et* al. (2015). Three days after transfection, culture media plus Y-27632 was exchanged with selection medium. After puromycin selection, surviving clones were picked and subjected to genotyping to detect the presence of homologous recombination.

For genotyping, genomic DNA was isolated using Viagen Direct PCR (Viagen). The presence of oncogenic and silent mutations in exons and insertions or deletions in introns of *BRAF, KRAS* and *NRAS* was verified by using the PCR product obtained using the following primers: KRAS^G12D/G13D^ exon fw 5’-GGCTCATTGCAACCTCGG-3’,

KRAS^G12D/G13D^ exon rv 5’-GTTGGCGCCTACCGGTGG-3’,

KRAS^G12D/G13D^ exon silent mutations fw 5’-GAGACGGAGTCTTGCTCTAT-3’,

KRAS^G12D/G13D^ exon silent mutations rv 5’-GCTGTATGGTTAACGCACTC-3’,

KRAS^G12D/G13D^ intron fw 5’-CCGCAGAACAGCAGTCTG-3’,

KRAS^G12D/G13D^ intron rv 5’-TGATGTCACAATACCAAG-3’,

NRAS^G12D^ exon fw 5’-CCGACTGATTACGTAGCG-3’,

NRAS^G12D^ exon rv 5’-GTTGGCGCCTACCGGTGG-3’,

NRAS^G12D^ exon silent mutations rv 5’-GGGATCATATTCATCCACG-3’,

NRAS^G12D^ intron fw 5’-CCGACTGATTACGTAGCG-3’,

NRAS^G12D^ intron rv 5’-CTCATGAATGAACTCAACAC-3’,

BRAF^V600E^ exon fw 5’-GGAGAGCAGGATACCACAGC-3’,

BRAF^V600E^ exon rv 5’-GTTGGCGCCTACCGGTGG-3’,

BRAF^V600E^ exon silent mutations rv 5’-AACGTGACTTCTCCGTGGCC-3’,

BRAF^V600E^ intron fw 5’-CTTCATAATGCTTGCTCTG-3’,

BRAF^V600E^ intron rv 5’-CCTGCCTTAAATTGCATAC-3’.

Products were sequenced using the following primers:

KRAS^G12D/G13D^_exon 5’-CACCGATACACGTCTGCAGTCAAC-3’,

NRAS^G12D^ exon 5’-CCAAATGGAAGGTCACAC-3’,

BRAF^V600E^ exon 5’-CTTCATAATGCTTGCTCTG-3’,

In addition, the CloneJET PCR Cloning Kit was used to confirm indel generation in introns of the allele of *BRAF, KRAS* and *NRAS* knock-ins #1 that was subject to NHEJ.

### Vector construction

The CRISPR guide RNA (sgRNAs) were designed by an online CRISPR design tool (http://crispr.mit.edu). The sgRNA guide sequences used can be found in the supplementary data (Supplementary Table 4). For CRISPR-mediated homologous recombination the human codon-optimized Cas9 expression plasmid was obtained from Addgene (41815). The sgRNA-GFP plasmid was obtained from Addgene (41819) and used as a template for generating target specific sgRNAs as described in detail by Drost *et al*. (2015). For the generation of the donor template, genomic DNA from P18T organoids was used to PCR amplify the *NRAS, KRAS* and *BRAF* 3’ homology arms using high-fidelity Phusion Polymerase (New England BioLabs). The 5’ homology arm of *KRAS* spans the region Chr12:25245385-25245994, the 3’ homology arm spans the region Chr12:25244479-25245229. The 5’ homology arm of *NRAS* spans the region Chr1:114716161-114716768, the 3’ homology arm spans the region Chr1:114715261-114716010. The 5’ homology arm of *BRAF* spans the region Chr7:140753392-140753973, the 3’ homology arm spans the region Chr7:140752465-140753206. Gene block fragments (idtDNA) were used to generate 5’ homology arms containing silent and oncogenic mutations. The gene block fragments and homology arms were cloned into a pBlueScript plasmid expressing a 3229-bp AATPB:PGKpuroDtk selection cassette (Schwank *et al*., 2013).

### Western blot assay and RAS-GTP pull down

Prior to cell lysis, organoids were incubated with 1 mg/ml dispase II (Invitrogen) for 10 minutes at 37° C to digest the BME. Western blot samples for phosphorylated ERK and MEK were lysed using RIPA buffer (50 mM Tris-HCL pH 8.0, 150 mM NaCl, 0.1% SDS, 0.5% Na-Deoxycholate, 1% NP-40) containing Complete protease inhibitors (Roche). Protein content was quantified using a BCA protein assay kit (Pierce™) and analyzed by Western blotting. Membranes were blocked and probed with antibodies directed against pMEK (RRID:AB_331648), MEK (RRID:AB_823567), pERK (RRID:AB_331646), ERK (RRID:AB_390779) and GAPDH (RRID:AB_2107445). Samples for RAS-GTP isolation were lysed using Ral lysis buffer (50 mM Tris-HCL pH 7.5, 200 mM NaCl, 2 mM MgCl2, 10% glycerol, 1% NP-40) containing Complete protease inhibitors (Roche). Lysates were normalized for protein levels using a BCA protein assay kit (Pierce™) and subsequently GTP-bound RAS was isolated via immunoprecipitation using recombinant RAS binding domain of RAF1 (RAF1-RBD). Protein lysates were run on SDS-PAGE gels and transferred to PVDF membranes (Millipore). Membranes were blocked and probed with antibodies directed against RAS (RRID:AB_397425) and NRAS (RRID:AB_628041). Organoid treatments: afatinib (Selleck Chemicals) 1 µM, 1 h and 24 h or DMSO.

### RNA isolation, cDNA preparation and qRT–PCR

Organoids were harvested in RLT lysis buffer and RNA was isolated using the Qiagen RNeasy kit (Qiagen) according to the manufacturer’s instructions. Extracted RNA was used as a template for cDNA production using iScript™ cDNA Synthesis Kit (Bio-Rad) according to the manufacturer’s protocol. qRT–PCR was performed using FastStart Universal SYBR Green Master mix (Roche) according to the manufacturer’s protocol. Results were calculated by using the relative standard curve method. Primer sequences:

B2M_fw 5’-GAGGCTATCCAGCGTACTCCA-3’,

B2M_rv 5’-CGGCAGGCATACTCATCTTTT-3’,

KRAS wild-type fw 5’-GCAAGAGTGCCTTGACGATAC-3’,

KRAS wild-type rv 5’-CTGCTGTGTCGAGAATATC-3’,

KRAS mutant fw 5’-GCAAGAGTGCGTTAACCATAC-3’,

KRAS mutant rv 5’-CTGCTGTGTCGAGAATATC-3’,

NRAS wild-type fw 5’-TCCAGCTAATCCAGAACCAC-3’,

NRAS wild-type rv 5’-CCAGCTGTATCCAGTATGTC-3’,

NRAS mutant fw 5’-GCGCACTGACAATCCAATTG-3’,

NRAS mutant rv 5’-CCAGCTGTATCCAGTATGTC-3’,

BRAF wild-type fw 5’-ATCTCGATGGAGTGGGTC-3’,

BRAF wild-type rv 5’-CTGGTCCCTGTTGTTGATG-3’,

BRAF mutant fw 5’-CACGGAGAAGTCACGTTG-3’,

BRAF mutant rv 5’-GGTAACTGTCCAGTCATC-3’,

EGFR fw 5’-AGTGCCTGAATACATAAACC-3’,

EGFR rv 5’-GTAGTGTGGGTCTCTGC-3’,

HER2 fw 5’-TGTGACTGCCTGTCCCTACAA-3’,

HER2 rv 5’-CCAGACCATAGCACACTCGG-3’,

HER3 fw 5’-ATACACACCTCAAAGGTACTC-3’,

HER3 rv 5’-ATCTTCTTCTTCAGTACCCAG-3’.

### Phenotypic drug screen and Calcein Green Assay

Five days after organoid trypsinization, 1 mg/ml dispase II (Invitrogen) was added to the medium of the organoids and these were incubated for 15 min at 37° C to digest the BME. Subsequently, organoids were mechanically dissociated by pipetting, filtrated using a 40 μm nylon cell strainer (Falcon), resuspended in 75% BME/growth medium (40 organoids/µl) prior plating of two 10 µl drops on Nunc™Lab-Tek™II Chamber Slide™Systems. After plating culture medium containing either 1 µM of afatinib, 1 µM of erlotinib or DMSO was added. The labtek plates were mounted on an inverted confocal laser scanning microscope (Leica SP8X) and imaged using a 10X objective. For visualization of cell viability, organoids were incubated with 16.2 µM Hoechst 33342 (Life Technologies) and 1.5 µM DRAQ7™(Cell Signaling #7406) for 30 min at 37° C prior imaging. For calculating organoid viability, the morphology of 100 organoids was scored after 3 and 7 days of 1 µM afatinib (pan-HERi) treatment.

For organoid viability and growth analysis, organoids were imaged by an inverted routine microscope (Nikon Eclipse TS100) using a 4X objective. For calculating organoid count and size, organoids were incubated for 20 minutes with 500 ml culture medium containing 5 µM calcein-green (Invitrogen). For the quantification of the organoid size and count, FIJI analysis software was used and presented as dot plots^74^.

### Drug screen and viability assessment

Five days after organoid trypsinization to single cells, 1 mg/ml dispase II (Invitrogen) was added to the medium of the organoids and these were incubated for 15 min at 37° C to digest the BME. Subsequently, organoids were mechanically dissociated from the BME by subtle pipetting, filtrated using a 40 µm nylon cell strainer (Falcon), resuspended in 2% BME/growth medium (15–20,000 organoids/ml) prior plating of 30 µl (72 hrs drug screen) or 50 µl (7 days drugscreen) (Multi-dropTM Combi Reagent Dispenser) on BME pre-coated 384-well plates. The drugs and their combinations were added 3 hrs after plating the organoids by using the Tecan D300e Digital Dispenser. Drugs were dispensed in a non-randomized manner and DMSO end concentration was 0.9% in all wells. 72 hrs or 7 days after adding the drugs organoids were fixed with 4% PFA (Merck) and stained with Hoechst (Invitrogen). Organoids were screened by automated microscopy of whole wells (CX5 High Content Screening (HCS) platform (Thermo Scientific), equipped with an Olympus UPLFLN U Plan Fluorite 4x Microscope Objective). Organoid roundness was measured by integrating Hoechst signal and contrast using Columbus Cellular imaging and analyses (Perkin Elmer). Relative survival was determined by normalization of the results to DMSO (= 100% alive) and 20 µM Navitoclax (= 0% alive), which induces maximal killing within 72 hours after treatment. Multiple identical drug combinations were averaged.

### Targeted inhibitors

Afatinib, Selumetinib, Vemurafenib, Erlotinib and Navitoclax were purchased from Selleck Chemicals. SCH772984 was obtained from MedChem Express. These compounds were dissolved in dimethylsulfoxide (DMSO, Sigma-Aldrich) and stored as 10 mM aliquots.

### Curve fitting and drug sensitivity

Dose-response curves were generated using GraphPad software by performing nonlinear regression (curve fit), assuming a standard Hill equation (chosen method: log(inhibitor) vs. Response, constrain top=100).

### RNA seq

Organoids were treated with DMSO or afatinib (1 µM) for 24 hours and organoids were lysed in RLT lysis+ buffer containing 1% beta-mercaptoethanol. RNA was extracted with the QiaSymphony SP kit (Qiagen) according to the manufacturer’s protocol. RNA-libraries were prepared for sequencing with the Illumina Truseq Stranded mRNA polyA kit and sequenced on the NextSeq500 platform (1×75bp, 20M reads per sample). Data were processed by the UBEC facility Illumina analysis pipeline (https://github.com/UMCUGenetics/RNASeq) by utilizing STAR (v2.4.2a) to map the reads. Samples were normalized for sequencing depth based on the sum of the read counts over all genes for each sample. Expressed genes were selected by excluding all genes where ≥ 3 samples had less than 10 reads. Principal Component Analysis (PCA), Euclidean Distance-based clustering and Differential Expression (DGEA) calculations were performed with the DESeq2 package^75^. P-adjusted (p_adjusted_) was calculated by multiplying the p-value with the number of genes (=expressed genes) tested. Unsupervised hierarchical clustering was performed on all differentially expressed genes.

DGEA was performed on untreated parental P18T lines vs. untreated KRAS^G12D^, KRAS^G13D^, NRAS^G12D^ and BRAF^V600E^ mutant lines, on treated parental P18T lines vs. treated KRAS^G12D^, KRAS^G13D^, NRAS^G12D^ and BRAF^V600E^ mutant lines, on untreated vs treated lines per mutation, or on treated KRAS^G12D^ and BRAF^V600E^ vs KRAS^G13D^ and NRAS^G12D^ mutant lines. For subsequent analysis, we set a cut-off threshold of a log_2_ 2 fold change of genes that were differentially expressed in all three (mutant) or four (P18T) monoclonal lines.

### CNAs

Genomic DNA was extracted from the parental P18T and RAS pathway mutant organoid lines using the QIAamp DNA Micro Kit (Qiagen) according to the manufacturer’s protocol. Approximately 100 ng genomic DNA of each sample was used for DNA library preparation (TruSeq Nano DNA library preparation kit, Illumina). Subsequently, libraries were sequenced on an Illumina NextSeq500 using 75-bp single-end sequencing. Raw sequencing data was aligned to human reference genome hg19/GRCh37 using Burrows-Wheeler Aligner mapping tool (BWA-MEM; Version 0.7.5a). Further data processing procedures are fully described at: https://github.com/UMCUGenetics/IAP. DNA copy number profiles were generated using Ginkgo as described by Garvin et al. (2015)^76^ (pipeline available at: https://github.com/robertaboukhalil/ginkgo). The reads were binned into 1 Mb variable-length intervals and data was segmented to obtain copy number estimates across the genome. Copy number values deviating >0.6 from the average ploidy were considered to indicate deletions or amplifications. R package ggplot2 was used to generate a heatmap for visualization.

### Organoid xenograft experiments

Approval for this study was obtained by the local animal experimental committee at The Netherlands Cancer Institute (IVD-NKI; OZP 80102; WP8520). Parental P18T and RAS pathway mutant knock-in patient-derived organoids were transplanted subcutaneous as single cells at a density of 3×10^5 organoids in 100 µl 50% Matrigel/medium with 10% collagen type I (BD Bio-sciences) mixture into NSG-B2m mice (JAX stock no: 010636).

Tumor growth dynamics were analyzed for 35 days in mice with established tumors of 50 mm^3^. Mice with established tumors (average volume of 200 mm^3^) were treated with afatinib (20 mg/kg; five days on, two days off) or vehicle for four weeks. After two weeks recovery from the drug treatment, mice were sacrificed.

Tumor volumes were evaluated three times per week by caliper and the approximate volume of the mass was calculated using the formula Dxd2/2, where D is the major tumor axis and d is the minor tumor axis. For in vivo dosing, afatinib was dissolved in 1.8% hydroxypropyl-b-cyclodextrin (Sigma), 5% of a 10% acetic acid stock and aqueous natrosol (0,5%). All agents were administered via oral gavage. We used 5 mice per group that were randomly assigned to the different treatment groups before the start of the experiment. We determined outliers with the following rule: If a number is less than Q1-1.5xIQR or greater than Q3+1.5xIQ, then it is considered to be an outlier, with IQR being the interquartile range, equal to the difference between the third quartile (Q3) and first quartile (Q1). Mice that showed outliers in more than 40% of the total number of measurements were excluded from analysis.

### Statistical analysis

GraphPad Prism 8.1.1 was used for statistical analysis. All values are given as means ± SD, as indicated in figure legends. Comparison between two groups were made by Welch’s *t-*test. For comparison of more than two groups, we used 2-way ANOVA with subsequent Dunnett’s or Bonferroni’s multiple comparison test.

## Authors’ Contributions

JBP and HJGS conceived the study. JBP, HJGS designed experiments, and JBP performed most of the experiments. JBP generated clones. NH performed drug screens. JBP and NH analyzed and interpreted the data. IV cultured and characterized CRC PDOs by targeted genome sequencing. JL, MV and RK performed in vivo mouse experiments, which was analyzed and interpreted by JBP. CS performed RNA sequencing analysis. JBP and CS analyzed and interpreted the data. ES performed and analyzed CNA experiments, which was interpreted by JBP and HJGS. JBP and HJGS wrote the manuscript, which was reviewed by all authors.

## Acknowledgements

This work is part of the Oncode Institute, which is partly financed by the Dutch Cancer Society, and was funded by the gravitation program CancerGenomiCs.nl from the Netherlands Organization for Scientific Research (NWO), by a grant from the Dutch Cancer Society (UU 2013-6070), by a ‘Sta op tegen Kanker’ International Translational Cancer Research Grant, and ERC starting grant (H.J.G.S). Stand Up to Cancer is a program administered by the AACR. JvR was supported by an ERC CoG Cancer-recurrence grant (648804), Cancer Genomics Netherlands and the Doctor Josef Steiner Foundation. Furthermore, we thank all members of the Snippert, Gloerich and De Rooij laboratories for fruitful discussions and support. We thank Utrecht Sequencing Facility for providing sequencing service and data. Utrecht Sequencing Facility is subsidized by the University Medical Center Utrecht, Hubrecht Institute,Utrecht University and The Netherlands X-omics Initiative (NWO). We thank the UMCU Bioinformatics Expertise Core (UBEC) facility for their help with the analysis of RNA sequencing results. Last, we would like to thank the people from the Preclinical Intervention Unit of the Mouse Clinic for Cancer and Ageing (MCCA) at the NKI for performing the intervention studies.

