## Supplementary Figures for "Cancer modeling in colorectal organoids reveals intrinsic differences between oncogenic *RAS* and *BRAF* variants"

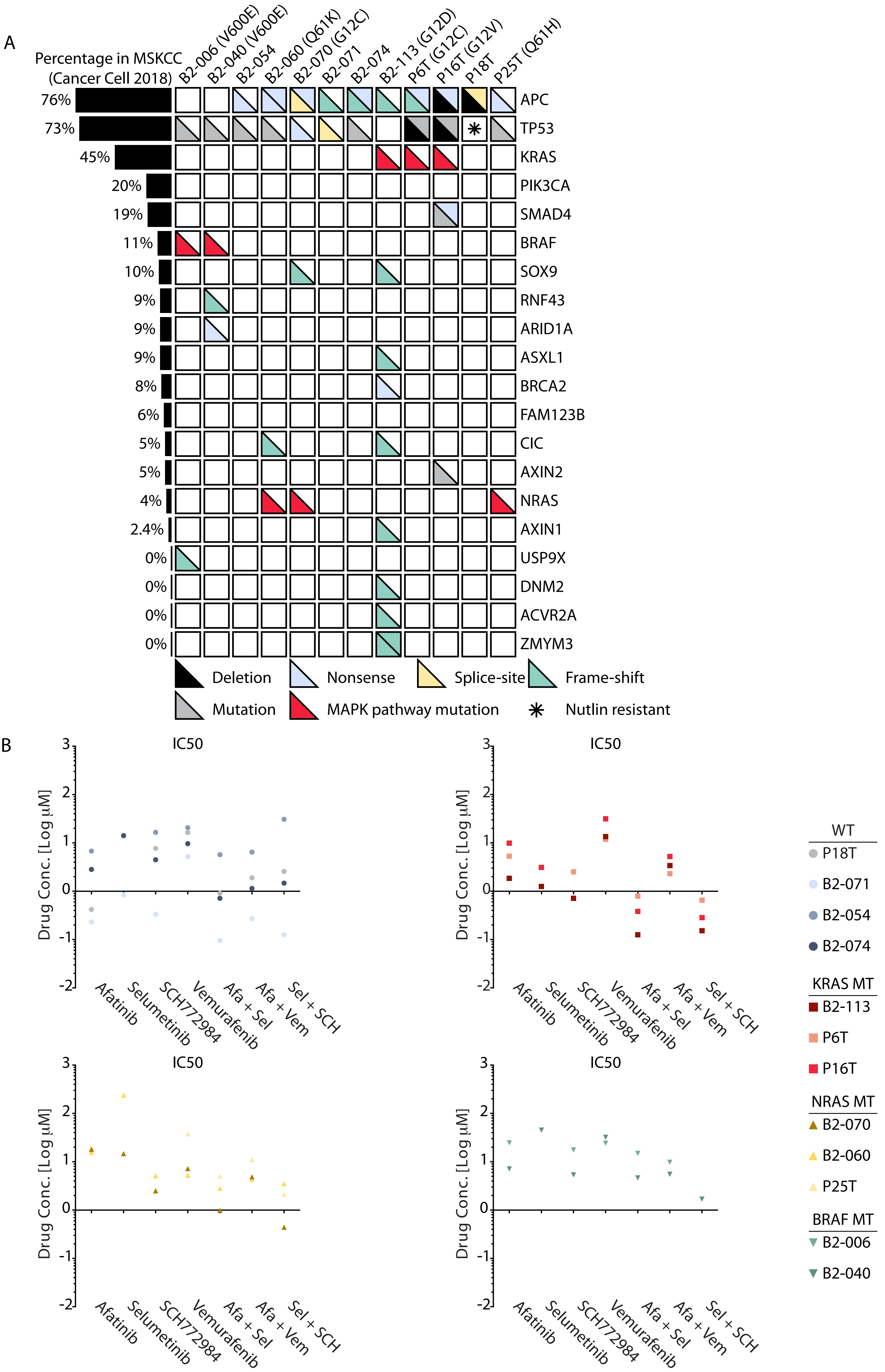
A
Percentage in MSKCC (Cancer Cell 2018)
B2-006 (V600E)
B2-040 (V600E)
B2-054
B2-060 (Q61K)
B2-070 (G12C)
B2-071
B2-074
B2-113 (G12D)
P6T (G12C)
P16T (G12V)
P18T
P25T (Q61H)
76% APC
73% TP53
45% KRAS
20% PIK3CA
19% SMAD4
11% BRAF
10% SOX9
9% RNF43
9% ARID1A
9% ASXL1
8% BRCA2
6% FAM123B
5% CIC
5% AXIN2
4% NRAS
2.4% AXIN1
0% USP9X
0% DNM2
0% ACVR2A
0% ZMYM3
Deletion
Nonsense
Splice-site
Frame-shift
Mutation
MAPK pathway mutation
Nutlin resistant
B
IC50
Drug Conc. [Log µM]
Afatinib
Selumetinib
SCH772984
Vemurafenib
Afa + Sel
Afa + Vem
Sel + SCH
WT
P18T
B2-071
B2-054
B2-074
KRAS MT
B2-113
P6T
P16T
NRAS MT
B2-070
B2-060
P25T
BRAF MT
B2-006
B2-040

**Supplementary Figure 1. Overview of common cancer mutations in selected CRC PDOs with oncogenic mutant MAPK signaling.**

(**A**) Overview of common cancer mutations identified in the selected CRC PDOs with oncogenic mutations in the MAPK pathway. Each box (per half) represents both alleles and indicates a deletion, missense mutation, nonsense mutation, splice-site mutation or a frame-shift alteration according to legend. MAPK pathway mutations are marked in red. The asterisk in P18T indicates a non-functional TP53 pathway (e.g. Nutlin-3 resistant). Genetic alterations in the panel of CRC PDOs are compared to the frequency of mutations as reported in clinical samples (cBioPortal) and are indicated on the left. CRC PDOs identities is indicated at the top, gene names on the right. (**B**) IC50 values derived from dose-response measurements of CRC PDOs treated with afatinib, selumetinib, SCH772984 and vemurafenib in a mono- or combinatorial fashion (see **Fig. 1**). Panels are subdivided per MAPK mutation in the CRC PDO lines. Line identity is indicated in the legend.

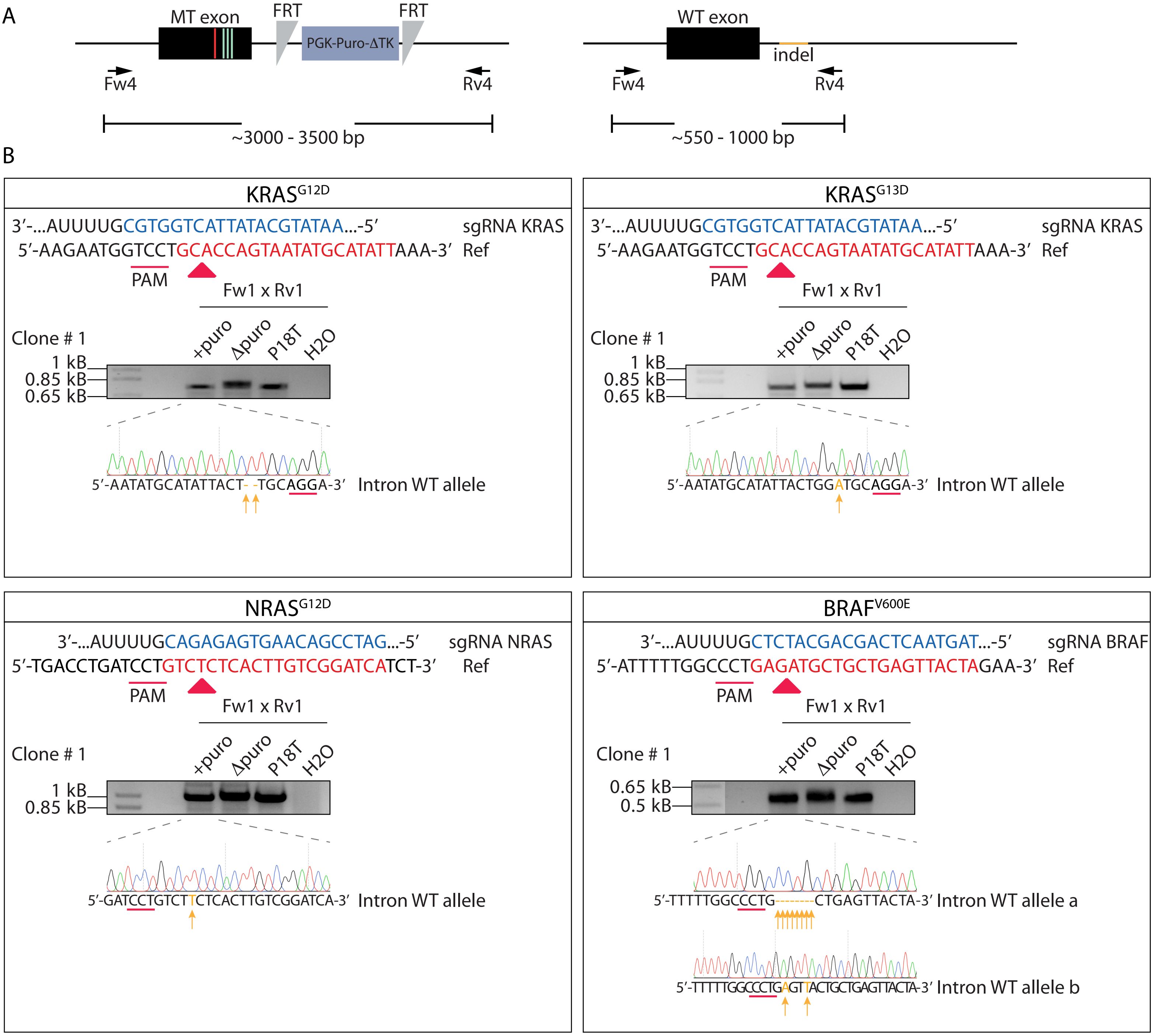
A
MT exon
FRT
PGK-Puro-ΔTK
FRT
Fw4
Rv4
~3000 - 3500 bp
WT exon
indel
Fw4
Rv4
~550 - 1000 bp
B
KRAS G12D
3′-...AUUUUGCGTGGTCATTATACGTATAA...-5′ sgRNA KRAS
5′-AAGAATGGTCCTGCACCAGTAATATGCATATTAAA-3′ Ref
PAM
Fw1 x Rv1
Clone # 1
+puro Δpuro P18T H2O
1 kB
0.85 kB
0.65 kB
5′-AATATGCATATTACT- -TGCAGGA-3′ Intron WT allele
KRAS G13D
3′-...AUUUUGCGTGGTCATTATACGTATAA...-5′ sgRNA KRAS
5′-AAGAATGGTCCTGCACCAGTAATATGCATATTAAA-3′ Ref
PAM
Fw1 x Rv1
Clone # 1
+puro Δpuro P18T H2O
1 kB
0.85 kB
0.65 kB
5′-AATATGCATATTACTGGATGCAGGA-3′ Intron WT allele
NRAS G12D
3′-...AUUUUGCAGAGAGTGAACAGCCTAG...-5′ sgRNA NRAS
5′-TGACCTGATCCTGTCTCTCACTTGTCGGATCATCT-3′ Ref
PAM
Fw1 x Rv1
Clone # 1
+puro Δpuro P18T H2O
1 kB
0.85 kB
5′-GATCCTGTCTTCTCACTTGTCGGATCA-3′ Intron WT allele
BRAF V600E
3′-...AUUUUGCTCTACGACGACTCAATGAT...-5′ sgRNA BRAF
5′-ATTTTTGGCCCTGAGATGCTGCTGAGTTACTAGAA-3′ Ref
PAM
Fw1 x Rv1
Clone # 1
+puro Δpuro P18T H2O
0.65 kB
0.5 kB
5′-TTTTTGGCCCTG--------CTGAGTTACTA-3′ Intron WT allele a
5′-TTTTTGGCCCTGAGTTACTGCTGAGTTACTA-3′ Intron WT allele b

**Supplementary Figure 2. Intronic indel mutations during generation of oncogenic *BRAF* and *RAS* knock-in variants in CRC PDOs**.

(**A**) Genetic strategy to identify the introduction of indel mutations in the introns of *KRAS*, *NRAS* and *BRAF* alleles as a result of Cas9 targeting. In the absence of homologous recombination, double strand breaks are repaired by NHEJ. Black arrows illustrate PCR primer pairs, which amplify genomic loci from both alleles that can be discriminated based on size. The smaller product that includes Cas9-targeted location (orange) was sent for Sanger sequencing (**B**) Per mutation the gRNA target site is shown, as well as the agarose electrophoresis gels showing the PCR product of the allele that was repaired by NHEJ (all data from clones # 1). Sanger sequencing indicates the introduction of small indel mutations in the intronic region. Nonmatching bases are shown in orange. Regions of the sgRNA complementary to the protospacer (underlined) are shown in blue. Red arrow heads indicate Cas9 cleavage sites. Of note, Sanger sequencing of clone # 1 of $BRAF^{V600E}$ mutant CRC PDOs shows the presence of multiple 'normal' alleles (>2N). +Puro indicates the monoclonal line that contains the puromycin selection cassette. ∆Puro indicates derivative of the monoclonal lines #1 in which the puromycin selection cassette was removed via FLP-FRT recombination.

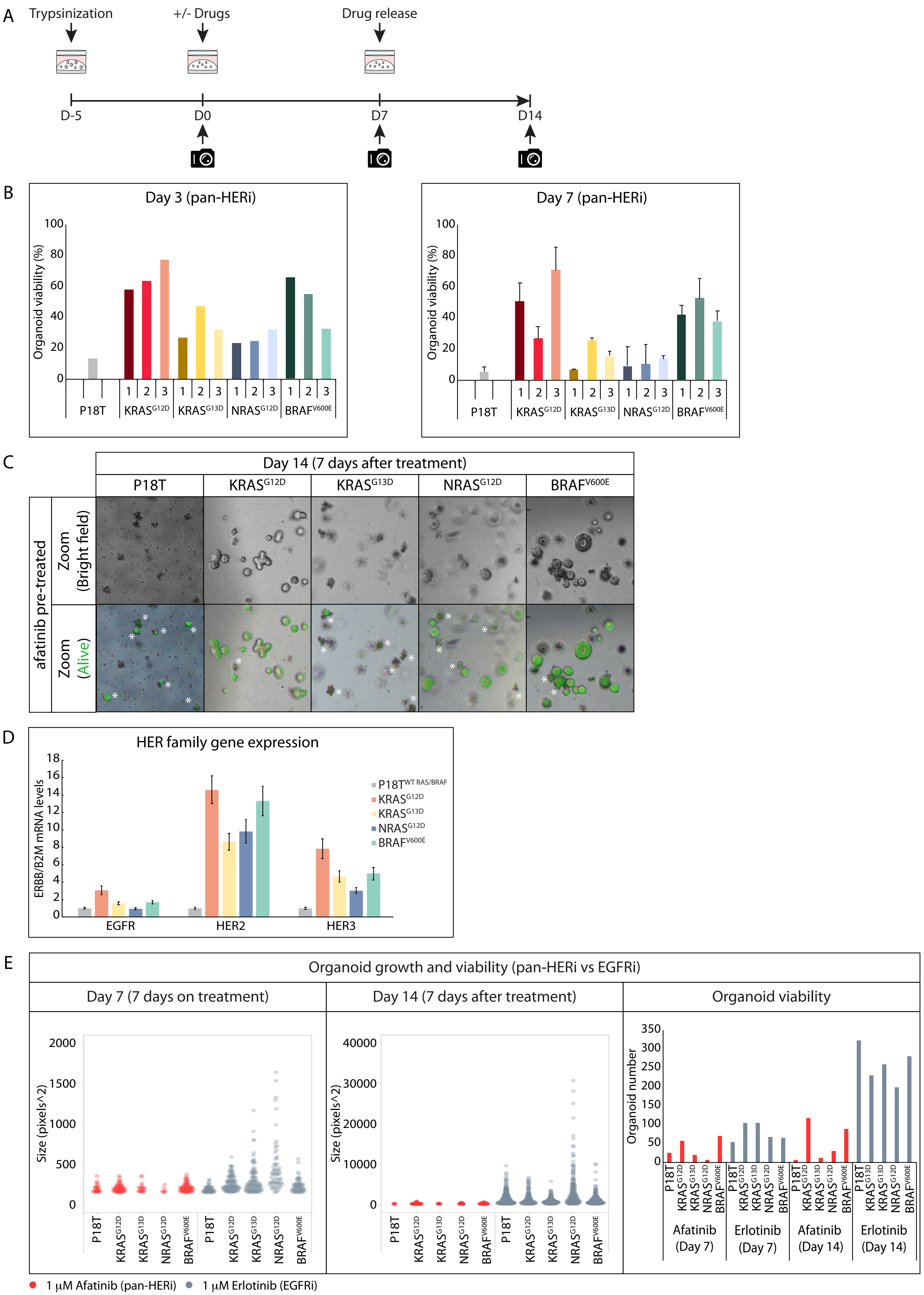

A
Trypsinization
+/- Drugs
Drug release
D-5
D0
D7
D14
B
Day 3 (pan-HERi)
Day 7 (pan-HERi)
Organoid viability (%)
P18T
KRAS G12D
KRAS G13D
NRAS G12D
BRAF V600E
C
Day 14 (7 days after treatment)
afatinib pre-treated
Zoom (Bright field)
Zoom (Alive)
D
HER family gene expression
ERBB/B2M mRNA levels
EGFR
HER2
HER3
P18T WT RAS/BRAF
E
Organoid growth and viability (pan-HERi vs EGFRi)
Day 7 (7 days on treatment)
Day 14 (7 days after treatment)
Organoid viability
Size (pixels^2)
Organoid number
Afatinib (Day 7)
Erlotinib (Day 7)
Afatinib (Day 14)
Erlotinib (Day 14)
1 µM Afatinib (pan-HERi)
1 µM Erlotinib (EGFRi)

**Supplementary Figure 3.** ***KRAS***$^{G12D}$ **and** ***BRAF***$^{V600E}$ **knock-in mutations promote organoid survival in the absence of EGFR signaling activity.**

(**A**) Schematic overview of the strategy to score viability and growth of parental P18T and *KRAS*$^{G12D}$, *KRAS*$^{G13D}$, *NRAS*$^{G12D}$ and *BRAF*$^{V600E}$ knock-in organoids. Growth from single cells ensures homogeneous size distribution during the experiment. Organoid size and viability were quantified after 3 and 7 days of drug treatment and 7 days after drug withdrawal using the selective uptake of fluorescent calcein green by viable cells. (**B**) Bar graphs depict the percentage of viable organoids after 3 and 7 days of pan-HER inhibition (1 μM afatinib). (**C**) Representative zoom-in panels of parental P18T, or *KRAS*$^{G12D}$, *KRAS*$^{G13D}$, *NRAS*$^{G12D}$ and *BRAF*$^{V600E}$ knock-in organoids (clones # 1) 7 days after release of afatinib treatment (day 14). Upper panel shows bright field pictures, lower panel shows overlay with fluorescent calcein green signal in living cells. Asterisks indicate autofluorescence of dead material. (**D**) mRNA expression levels of EGFR, HER2 and HER3 in parental P18T and *KRAS*$^{G12D}$, *KRAS*$^{G13D}$, *NRAS*$^{G12D}$ and *BRAF*$^{V600E}$ knock-in organoids measured by qPCR analysis, normalized over housekeeping gene *B2M* (n=3). (**E**) Quantitative analysis of organoid size and viability of isogenic RPM lines during (day 7) and after (day 14) treatment with pan-HER inhibition (1 μM afatinib, red dots) or EGFR selective inhibition (1 μM erlotinib, grey dots). Size and number of viable organoids were measured by selective uptake of fluorescent calcein green by viable cells (see methods). Each dot represents one organoid[76]. Data from 1 8-well Lab-Tek chambered coverglass is shown.

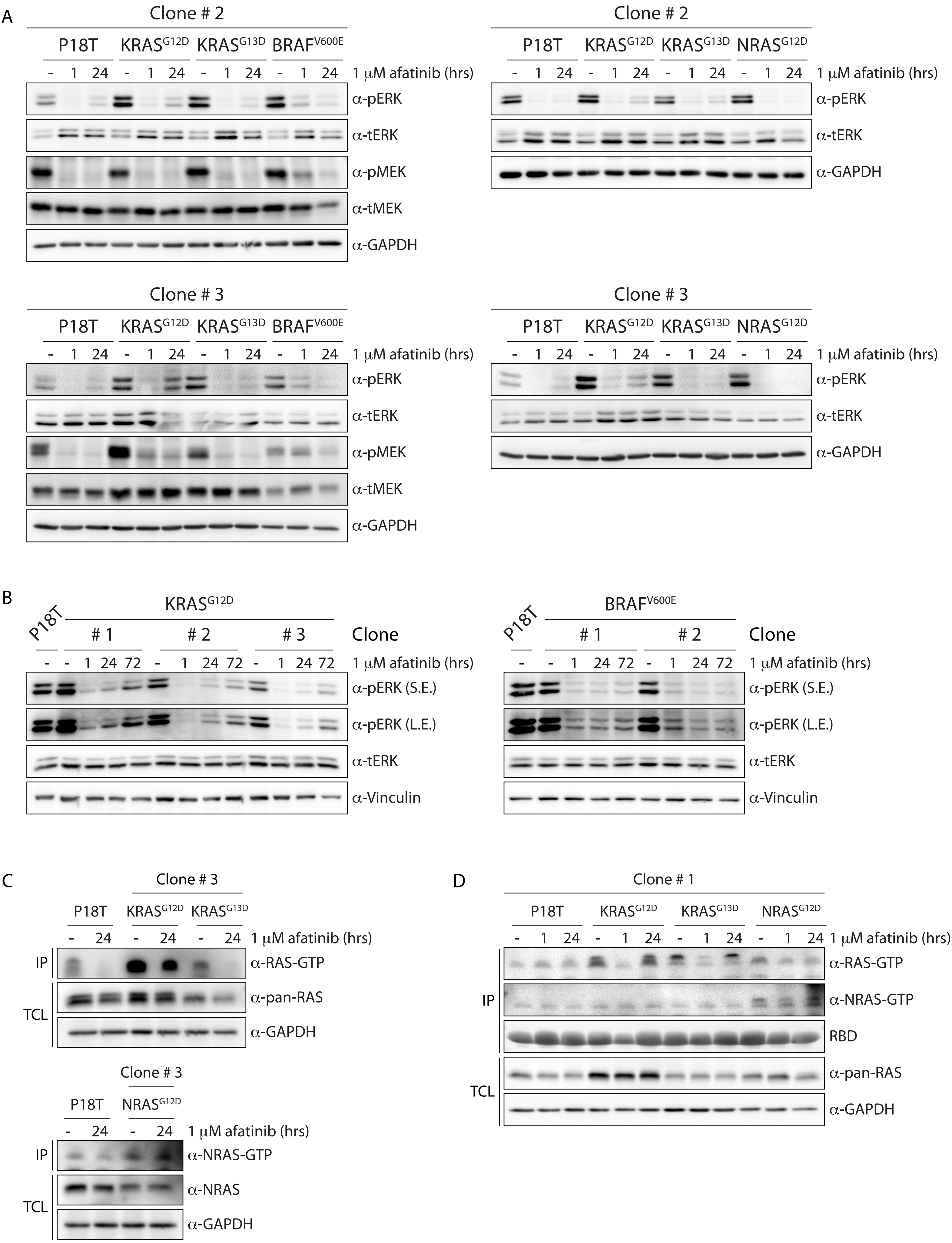

A
Clone # 2
P18T
KRAS$^{G12D}$
KRAS$^{G13D}$
BRAF$^{V600E}$
- 1 24 - 1 24 - 1 24 - 1 24
1 μM afatinib (hrs)
α-pERK
α-tERK
α-pMEK
α-tMEK
α-GAPDH
Clone # 2
P18T
KRAS$^{G12D}$
KRAS$^{G13D}$
NRAS$^{G12D}$
- 1 24 - 1 24 - 1 24 - 1 24
1 μM afatinib (hrs)
α-pERK
α-tERK
α-GAPDH
Clone # 3
P18T
KRAS$^{G12D}$
KRAS$^{G13D}$
BRAF$^{V600E}$
- 1 24 - 1 24 - 1 24 - 1 24
1 μM afatinib (hrs)
α-pERK
α-tERK
α-pMEK
α-tMEK
α-GAPDH
Clone # 3
P18T
KRAS$^{G12D}$
KRAS$^{G13D}$
NRAS$^{G12D}$
- 1 24 - 1 24 - 1 24 - 1 24
1 μM afatinib (hrs)
α-pERK
α-tERK
α-GAPDH
B
P18T
KRAS$^{G12D}$
# 1
# 2
# 3
Clone
- - 1 24 72 - 1 24 72 - 1 24 72
1 μM afatinib (hrs)
α-pERK (S.E.)
α-pERK (L.E.)
α-tERK
α-Vinculin
P18T
BRAF$^{V600E}$
# 1
# 2
Clone
- - 1 24 72 - 1 24 72
1 μM afatinib (hrs)
α-pERK (S.E.)
α-pERK (L.E.)
α-tERK
α-Vinculin
C
Clone # 3
P18T
KRAS$^{G12D}$
KRAS$^{G13D}$
- 24 - 24 - 24
1 μM afatinib (hrs)
IP
α-RAS-GTP
TCL
α-pan-RAS
α-GAPDH
Clone # 3
P18T
NRAS$^{G12D}$
- 24 - 24
1 μM afatinib (hrs)
IP
α-NRAS-GTP
TCL
α-NRAS
α-GAPDH
D
Clone # 1
P18T
KRAS$^{G12D}$
KRAS$^{G13D}$
NRAS$^{G12D}$
- 1 24 - 1 24 - 1 24 - 1 24
1 μM afatinib (hrs)
IP
α-RAS-GTP
α-NRAS-GTP
RBD
TCL
α-pan-RAS
α-GAPDH

**Supplementary Figure 4. *KRAS*$^{G12D}$ and *BRAF*$^{V600E}$ RPM lines show residual MAPK pathway activity in the presence of pan-HER inhibition.**

(**A**) Organoids expressing oncogenic *KRAS* (G12D and G13D), *NRAS* (G12D) and *BRAF* (V600E) variants show enhanced basal ERK phosphorylation levels compared to P18T organoids. Pan-HER inhibition (1 µM afatinib) shows sustained ERK and MEK phosphorylation in *KRAS*$^{G12D}$ and *BRAF*$^{V600E}$ organoids compared to P18T, *KRAS*$^{G13D}$ and *NRAS*$^{G12D}$ organoids. Top panels are representative biochemistry experiments on clones # 2 from n=3. Bottom panels are representative biochemistry experiments on clones # 3 from n=3. (**B**) *KRAS*$^{G12D}$ organoids show ERK reactivation upon prolonged pan-HER inhibition (72 hours of 1 µM afatinib), whereas low levels of ERK phosphorylation remain stable in *BRAF*$^{V600E}$ organoids. Left panel (KRAS$^{G12D}$) are representative biochemistry experiments on clones # 1-3 from n=1. Right panel (BRAF$^{V600E}$) are representative biochemistry experiments on clones # 1-2 from n=1. S.E. short exposure, L.E. long exposure. (**C**) Biochemistry on RAS activity (GTP-loading) in unperturbed culture conditions and pan-HER inhibition (afatinib 1 µM) for *KRAS* (G12D and G13D) and *NRAS* (G12D) mutant clones # 3 compared to P18T CRC organoids. RAS immunoblots from RAS pull-down assay are shown (RAS-GTP), together with a RAS immunoblot from total cell lysates as loading control. HRAS, KRAS, and NRAS isoforms are detected in mutant KRAS pull-down assays. NRAS isoforms are detected in mutant NRAS pull-down assays. Representative from n = 3 independent experiments. (**D**) as in C but for clones # 1.

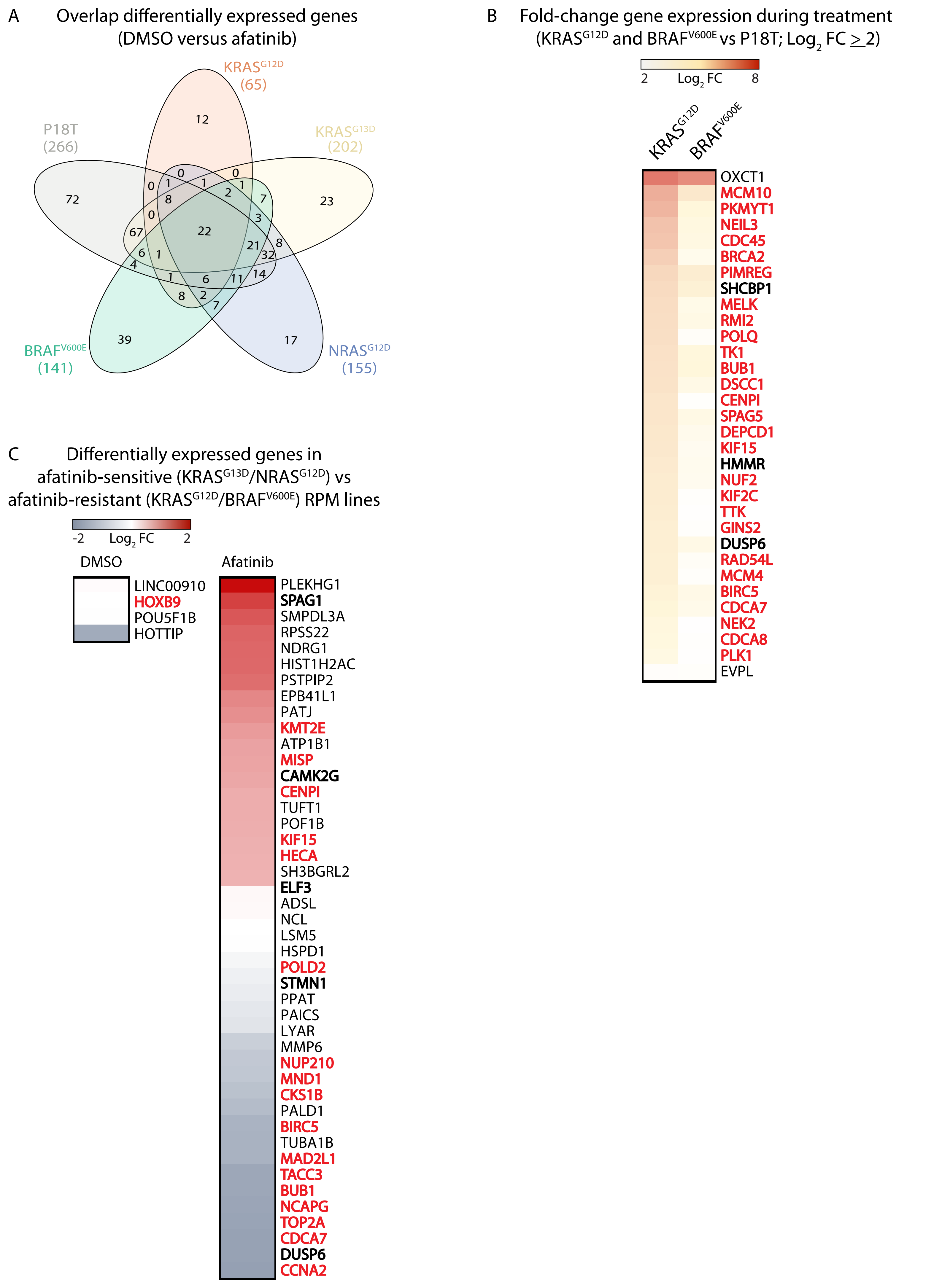
A
Overlap differentially expressed genes
(DMSO versus afatinib)
KRASG12D
(65)
KRASG13D
(202)
P18T
(266)
BRAFV600E
(141)
NRASG12D
(155)
12
72
23
39
17
0
1
1
0
0
1
8
2
7
0
3
67
22
21
8
6
1
32
4
14
1
6
11
8
2
7
B
Fold-change gene expression during treatment
(KRASG12D and BRAFV600E vs P18T; Log2 FC ≥ 2)
2
Log2 FC
8
KRASG12D
BRAFV600E
OXCT1
MCM10
PKMYT1
NEIL3
CDC45
BRCA2
PIMREG
SHCBP1
MELK
RMI2
POLQ
TK1
BUB1
DSCC1
CENPI
SPAG5
DEPCD1
KIF15
HMMR
NUF2
KIF2C
TTK
GINS2
DUSP6
RAD54L
MCM4
BIRC5
CDCA7
NEK2
CDCA8
PLK1
EVPL
C
Differentially expressed genes in
afatinib-sensitive (KRASG13D/NRASG12D) vs
afatinib-resistant (KRASG12D/BRAFV600E) RPM lines
-2
Log2 FC
2
DMSO
LINC00910
HOXB9
POU5F1B
HOTTIP
Afatinib
PLEKHG1
SPAG1
SMPDL3A
RPSS22
NDRG1
HIST1H2AC
PSTPIP2
EPB41L1
PATJ
KMT2E
ATP1B1
MISP
CAMK2G
CENPI
TUFT1
POF1B
KIF15
HECA
SH3BGRL2
ELF3
ADSL
NCL
LSM5
HSPD1
POLD2
STMN1
PPAT
PAICS
LYAR
MMP6
NUP210
MND1
CKS1B
PALD1
BIRC5
TUBA1B
MAD2L1
TACC3
BUB1
NCAPG
TOP2A
CDCA7
DUSP6
CCNA2

**Supplementary Figure 5. Gene expression profiles of RPM lines during unperturbed and EGF-independent growth.**

(**A**) Overlap of significantly downregulated genes ($p_{adjusted}$ <0.05; $log_2$ fold change (FC) $\leq$-2) upon pan-HER inhibition (1 µM afatinib) treatment versus DMSO in parental P18T (average of bulk and clones # 1-3) and $KRAS^{G12D}$, $KRAS^{G13D}$, $NRAS^{G12D}$ and $BRAF^{V600E}$ knock-in organoids (average of clones # 1-3). The EGFR gene signature in P18T of 266 genes includes 22 genes that are uniform targets in all lines. (**B**) Heatmaps of the genes (not-restricted to EGFR gene signature in P18T) with significantly higher expression ($p_{adjusted}$ <0.05; $log_2$ fold change (FC) $\geq$2) in $KRAS^{G12D}$ and $BRAF^{V600E}$ RPM lines when compared to parental P18T lines in treated (afatinib) conditions. (**C**) Heatmaps of the genes (not-restricted to EGFR gene signature in P18T) with differential expression ($p_{adjusted}$ <0.05; $log_2$ fold change (FC) $\geq$2) in afatinib-sensitive ($KRAS^{G13D}$ and $NRAS^{G12D}$) versus afatinib-resistant ($KRAS^{G12D}$ and $BRAF^{V600E}$) RPM lines during unperturbed (DMSO) and treated (1 µM afatinib) conditions. RAS-ERK pathway genes are depicted in bold. Cell cycle-related genes are marked in bold, red font.

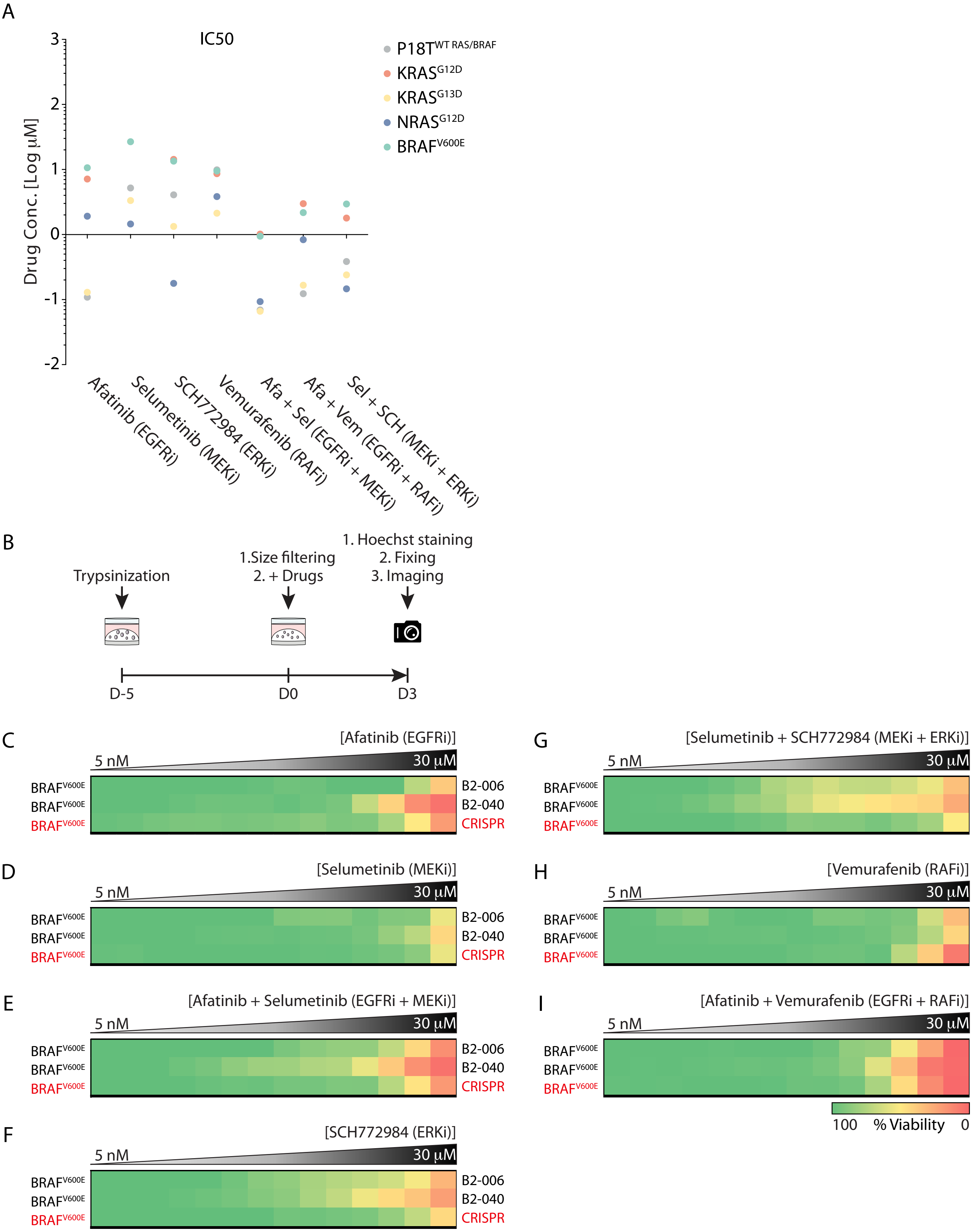
A
IC50
Drug Conc. [Log µM]
P18T WT RAS/BRAF
KRAS G12D
KRAS G13D
NRAS G12D
BRAF V600E
Afatinib (EGFRi)
Selumetinib (MEKi)
SCH772984 (ERKi)
Vemurafenib (RAFi)
Afa + Sel (EGFRi + MEKi)
Afa + Vem (EGFRi + RAFi)
Sel + SCH (MEKi + ERKi)
B
Trypsinization
1.Size filtering
2. + Drugs
1. Hoechst staining
2. Fixing
3. Imaging
D-5
D0
D3
C
[Afatinib (EGFRi)]
D
[Selumetinib (MEKi)]
E
[Afatinib + Selumetinib (EGFRi + MEKi)]
F
[SCH772984 (ERKi)]
G
[Selumetinib + SCH772984 (MEKi + ERKi)]
H
[Vemurafenib (RAFi)]
I
[Afatinib + Vemurafenib (EGFRi + RAFi)]
5 nM
30 µM
BRAF V600E
B2-006
B2-040
CRISPR
100
% Viability
0

**Supplementary Figure 6. Differential drug sensitivities of engineered RPM lines and patient-derived *BRAF*$^{V600E}$ CRC PDOs to targeted MAPK pathway inhibition.**

(**A**) IC50 values derived from dose-response measurements of RPM organoids treated with afatinib, selumetinib, SCH772984 and vemurafenib in a mono- or combinatorial fashion (see Fig. 6). Line identity is indicated in the legend. (**B**) Schematic overview of the 3 day drug screening method to compare CRC PDO lines with identical oncogenic *BRAF* mutation. (**C**) Heatmaps of drug response (viability) to afatinib, (**D**) selumetinib, (**E**) afatinib plus selumetinib, (**F**) SCH772984, (**G**) SCH772984 plus selumetinib, (**H**) vemurafenib and (**I**) vemurafenib plus afatinib. Organoids were treated (72 hr) with vehicle (DMSO) or inhibitors targeting the EGFR-RAS-ERK pathway (5 nM – 30 µM range, in 14 logarithmic intervals). Red represents maximal cell death and green represents maximal viability. Drug names and their nominal targets are indicated above, the MAPK mutant status per line at the left and the corresponding PDO line at the right. RPM organoids are indicated in red font. Average of 2 technical replicates.
